# Relating Global and Local Connectome Changes to Dementia and Targeted Gene Expression in Alzheimer’s Disease

**DOI:** 10.1101/730416

**Authors:** Samar S. M. Elsheikh, Emile R. Chimusa, Alzheimer’s Disease Neuroimaging Initiative, Nicola J. Mulder, Alessandro Crimi

## Abstract

Networks are present in many aspects of our lives, and networks in neuroscience have recently gained much attention leading to novel representations of brain connectivity. The integration of neuroimaging characteristics and genetics data allows a better understanding of the effects of the genetic variations on brain structural and functional connections. The current work uses whole-brain tractography in a longitudinal setting, and by measuring the brain structural connectivity changes studies the neurodegeneration of Alzheimer’s disease. This is accomplished by examining the effect of targeted genetic risk factors on the most common local and global brain connectivity measures. Furthermore, we examined the extent to which Clinical Dementia Rating relates to brain connections longitudinally, as well as to gene expression. Here we show that the expression of *PLAU* gene increases the change over time in betweenness centrality related to the fusiform gyrus. We also show that the betweenness centrality metric impact dementia-related changes in distinct brain regions. Our findings provide insights into the complex longitudinal interplay between genetics and brain characteristics and highlight the role of Alzheimer’s genetic risk factors in the estimation of regional brain connectivity alterations.

## Introduction

The advancement in technologies and the integration of genetic and neuroimaging datasets have taken Alzheimer’s research steps further, and produced detailed descriptions of molecular and brain aspects of the disease^1^. Previous studies have utilised the *connectome*^2^ *to study different brain diseases through associating genetic variants to brain connectivity*^*3*^. *A structural connectome* is a representation of the brain as a network of distinct brain regions (nodes) and their structural connections (edges), calculated as the number of anatomical tracts. Those anatomical tracts are generally obtained by diffusion-weighted imaging (DWI)^4^. DWI is the most commonly used in-vivo method for mapping and characterizing the diffusion of water molecules in three-dimensions, as a function of the location, and ultimately to construct the structural connectome. The connectome representation of the brain allows measuring of important properties, such as the ability of the brain to form separated sub-networks (*network segregation*), or the gathering of edges around specific nodes (*network integration*)^5^. Given such measures of the brain, it is also possible to represent each individual brain as single scalar metrics which summarize peculiar properties of the network’s segregation and integration^6^, and calculate what is known as global connectivity metrics. Moreover, these global metrics can also be used to quantify local properties at specific nodes/areas. This is relevant for neurodegeneration, as neuronal apoptosis in the end also reflects reduction of structural connectivity^7,8^.

There are many factors which may affect susceptibility to Alzheimer’s disease (AD) and various ways to measure the disease status. However, there is no single factor which can be used to predict the disease risk sufficiently^9^. Genetics is believed to be the most common risk factor in AD development^10^. Genetic variants located in about 20 genes have been reported to affect the disease through many cell-type specific biological functions^11^. Genome-Wide Associations Studies (GWAS), also highlighted dozens of multi-scale genetic variations associated with AD risk^8^,12,13. From the early stages of studying the disease, the well known genetic risk factors of AD were found to lie within the coding genes of proteins involved in amyloid-*β* (*Aβ*) processing. These include the Apolipoprotein E gene (*ApoE*) which increases the risk of developing AD^14^, the Amyloid precursor protein (APP)^15^, presenilin-1 (PSEN1) and presenilin-2 (PSEN2)^16,17^.

Early work demonstrated that *ApoE*-4 carriers have an accelerated age-related loss of global brain inter-connectivity in AD subjects^18^, and topological alterations of both structural and functional brain networks are present even in healthy subjects carrying the *ApoE* gene^19^. A meta-analysis study also showed the impact of APOE, phosphatidylinositol binding clathrin assembly protein (PICALM), clusterin (CLU), and bridging integrator 1 (BIN1) gene expression on resting state functional connectivity in AD patients^20^. Going beyond the *ApoE* gene, Jahanshad et al.^21^ used a dataset from the Alzheimer’s Disease Neuroimaging Initiative (ADNI) to carry out a GWAS of brain connectivity measures and found an associated variant in F-spondin (*SPON1*), previously known for its association with dementia severity. A more recent study has also used the ADNI dataset to conduct a GWAS on the longitudinal brain connectivity, and provided an evidence of association between some *single nucleotide polymorphisms* (SNPs) in the *CDH18* gene and Louvain modularity (a measure of network segregation)^8^.

AD is a common dementia-related illness; in the elderly, AD represents the most progressive and common form of dementia. Accordingly, incorporating and assessing dementia severity when studying AD provides more insights into the disease progression from a clinical point of view. A reliable global rating of dementia severity is the Clinical Dementia Rating (CDR)^22^, which represents a series of specific evaluations for memory, orientation, judgment and problem solving, community affairs, home and hobbies (intellectual interests maintained at home), and personal care. The ultimate CDR measure is an ordinal scale which rates the severity of dementia symptoms, it uses the values 0, 0.5, 1, 2 and 3, to represent none, very mild, mild, moderate and severe, respectively.

In this paper, we integrated different aspects of Alzheimer’s Disease, including longitudinal measures of CDR, global and local connectivity as well as gene expression extracted from blood samples of AD patients and controls. For global connectivity, we utilised the most commonly used metrics of network segregation and integration, such as the path length which was used in a plethora of studies as a biomarker to study the complex brain connectivity in schizophrenia^23^, and the local efficiency and characteristic path lengths that were used to study the structural organisation of the brain network in autism^24^. For a detailed review on work using such metrics see^25^. A brief description of the global connectivity metrics we used here, 1) *Louvain modularity* is a community (cluster) detection method, which iteratively transforms the network into a set of communities, 2) *transitivity* quantifies the segregation of a network by normalizing the fraction of triangles around an individual node, 3) *characteristic path length* is the average shortest path, 4) the *global efficiency* is the inverse of characteristic path length. For the local connectivity features, we used one measure to represent node’s segregation, integration and centrality. The local connectivity metrics of a network represent large-scale organization which in turn may be used to represent well functioning cognitive functions^26^. Measures of centrality (e.g. degree and betweeness centrality) usually measure similar features in a network, and hence, can be highly related to each other. Indeed, it has been shown that many of them can lead back to the degree centrality at a node^27^. Therefore, along with using local connectivity measures, we computed their correlation with the nodal degree metric -the simplest feature we can extract from a graph node.

Our work aimed to answer questions such as, are there brain connectivity metrics that discriminate changes longitudinally in AD patients compared to healthy control subjects? Is there a most representative metric, or a redundancy in the chosen metrics? Is there a correlation between the metrics used and expression of known AD-related genes? Is there a correlation between connectivity metrics and clinical ratings?

We address these questions considering the global brain by using global connectivity, and also considering specific brain regions using the local connectivity metrics. We regressed the absolute changes in connectivity metrics on gene expression, studied the association between the longitudinal changes of connectivity and CDR scores, and performed a ridge regression between gene expression, the changes in CDR scores and brain connectivity.

Using a dataset from ADNI (http://adni.loni.usc.edu/). this paper presents an integrated association study of specific AD risk genes, dementia scores and structural connectome characteristics. Specifically, we used the longitudinal case-control dataset to examine the association of known AD risk-gene expression with local and global connectivity metrics. Furthermore, we tested the longitudinal effect of brain connectivity on different CDR scores, and carried out a multivariate analysis to study the longitudinal effect of gene expression and connectome changes on CDR. Identifying advanced imaging and genetic bio-markers with regional effect is relevant in the clinical context as current efforts focus on treatments in specific locations and circuitry of the brain particularly affected by plaques^28^.

## Materials and Methods

### Data Description

We used two sets of data from ADNI, which are available at adni.loni.usc.edu. The experiments have been conducted on the publicly available datasets described below, for which ethical approval has already been granted, and data acquisition has been conducted according to the Helsinki II regulations. Additionally, we received ethics approval from the Faculty of Health Sciences Human Research Ethics Committee at the University of Cape Town.

The populations are matched by age, and the mean ages are respectively 76.5±7.4 for AD patients, and 77.0± 5.1 years for healthy subjects. To fulfil our objectives, unless otherwise specified, we merged neuroimaging, gene expression and CDR datasets for all the participants, who have these three types of data available, at two-time points. We considered follow-up imaging and CDR acquisition one year later than the baseline visit. All those constraints drastically reduced the number of available samples, yielding a total of 51 participants, 15 AD patients, and 36 healthy elderly subject. Recent studies^29^ focused on genetics and MRI data also used similar sample sizes. Therefore, despite this limitation this kind of study offers novel insights combining all those multimodal data generally not reached by purely imaging or non-longitudinal studies.

#### Imaging Data

For the imaging, we obtained the DWI volumes at two time points, the baseline and follow-up visits, with one year in between. Along with the DWI, we used the T1-weighted images which were acquired using a GE Signa scanner 3T (General Electric, Milwaukee, WI, USA). The T1-weighted scans were obtained with voxel size = 1.2×1.0 ×1.0*mm*^3^*TR* = 6.984*ms*; TE = 2.848 ms; flip angle= 11°, while DWI was obtained with voxel size = 1.4 ×1.4 ×2.7*mm*^3^, scan time = 9 min, and 46 volumes (5 T2-weighted images with no diffusion sensitization b0 and 41 diffusion-weighted images b= 1000*s/mm*^2^).

### Pre-processing of Imaging Data

Each DWI and T1 volume had been pre-processed performing eddy current correction and skull stripping. DWI and T1 volumes were already co-registered, and the atlas was further linearly registered to them according to 12 degrees of freedom.

We used the same T1 reference to get the information needed to compute the partial volume effect from the tissue segmentation by using the FMRIB Software Library (FSL). Non-linear registration was not necessary as volumes were registered according to individual patients, after the empirical observation that brain atrophy between the two time points was not pronounced enough to require non-linear methods. Although atrophy was visible mostly in the cingolum, it was not case for the cortex.

#### Genetic Data Acquisition

We used the Affymetrix Human Genome U219 Array profiled expression dataset from ADNI. The RNA was obtained from blood samples and normalised before hybridization to the array plates. Partek Genomic Suite 6.6 and Affymetrix Expression Console were used to check the quality of expression and hybridization^30^. The expression values were normalised using the Robust Multi-chip Average^31^, after which the probe sets were mapped according to the human genome (hg19). Further quality control steps were performed by checking the gender using specific gene expression, and predicting the SNPs from the expression data^32,33^. Although it is more useful to extract the gene expression profiles from the brain, we used the gene expression extracted from blood samples, as provided by ADNI. Blood gives a general idea of what is happening in the body, and can detect differences in gene expression. Moreover, blood samples are easy to obtain and are noninvasive. Nevertheless, we verified that the investigated genes are expressed in the brain parenchyma through the Allen gene expression atlas https://human.brain-map.org/ish/search. In this work, we targeted specific genes which have been reported to affect the susceptibility of AD. We used the BioMart software from Ensembl to choose those genes by specifying the phenotype as AD^34^. We obtained a total of 17 unique gene names and retrieved a total of 65 probe sets from the genetic dataset.

#### Clinical Dementia Rating

The Clinical Dementia Rating (CDR) score is an ordinal scale used to rate the condition of dementia symptoms. It ranges from 0 to 3, and is defined by five values: 0, 0.5, 1, 2 and 3, ordered by severity (smaller values are less severe), these values stand for none, very mild, mild, moderate and severe, respectively. The scores evaluate the cognitive state and functionality of participants. Here, we used the main six scores of CDR; memory, orientation, judgement and problem solving, community affairs, home and hobbies, and personal care. Besides these, we used a global score, calculated as the sum of the six scores. We obtained the CDR scores at two time points in accordance with the connectivity metrics time points.

#### Connectome Construction

We generated tractographies by processing DWI data with Dipy^35^, a Python library. More specifically, we used the constant solid angle model^36^ and Euler Delta Crossings^35^ deterministic algorithm. We stemmed from 2,000,000 seed-points, and as stopping condition we used the anatomically-constrained tractography approach based on partial volume effect^37^. We also discarded all tracts with sharp angle (larger than 75°) or those with length < 30*mm*.

To construct the connectome, we assigned a binary representation in the form of a matrix whenever more than three connections were present between two Automated anatomical labelling (AAL) atlas^38^ regions, for any pair of regions. Tracts shorter than 30 mm were discarded. Though the AAL atlas has been criticized for functional connectivity studies^39^, it has been useful in providing insights in neuroscience and physiology, and is believed to be sufficient for our case study limited to structural data and large brain regions^39^.

### Global and Local Connectivity Metrics

To quantify the overall efficiency and integrity of the brain, we extracted global measures of connectivity from the connectome, represented here in four values of network integration and segregation. Specifically, we used two network integration metrics 1) global efficiency (*E*), and 2) weighted characteristic path length (*L*). Both are used to measure the efficiency at which information is circulated around a network. On the other hand, we used; 1) Louvain modularity (*Q*), and, 2) transitivity (*T*) to measure the segregation of the brain which is defined as the capability of the network to shape sub-communities which are loosely connected to one another while forming a densely connected sub-network within communities^5,6^.

Suppose that *n* is the number of nodes in the network, *N* is the set of all nodes, the link (*i, j*) connects node *i* with node *j* and *a*_*i j*_ define the connection status between node *i* and *j*, such that *a*_*i j*_ = 1 if the link (*i, j*) exist, and *a*_*i j*_ = 0 otherwise. We define the global connectivity metrics as; 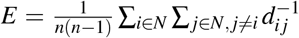, where,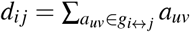, is the shortest path length between node *i* and *j*, and *g*_*i* ↔ *j*_ is the geodesic between *i* and *j*.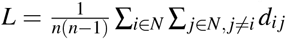. 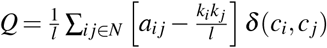, where *l* = ∑_*i, j*_*∈*_*N*_ *a*_*i j*_, *m*_*i*_ and *m*_*j*_ are the modules containing node *i* and *j*, respectively, and *δ* (*c*_*i*_, *c* _*j*_) = 1 if *c*_*i*_ = *c* _*j*_ and 0 otherwise. 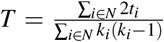, where 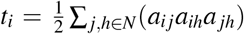 is the number of triangles around node *i*.

Moreover, using the AAL atlas, we constructed the following local brain network metrics at each region or node. We used the local efficiency (*E*_*loc,i*_), clustering coefficient (*C*_*i*_) and betweenness centrality (*b*_*i*_) at each node to quantify the local connectivity. Local efficiency and clustering coefficient measure the presence of well-connected clusters around the node, and they are highly correlated to each other. The betweenness centrality is the number of shortest paths which pass through the node, and measures the effect of the node on the overall flow of information in the network^6^. The local connectivity metrics used in this work, for a single node *i*, are defined as follows;

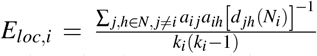, where, *d* _*jh*_(*N*_*i*_), is the length of the shortest path between node *j* and *h* - as defined in Equation, and contains only neighbours of h 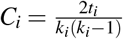. 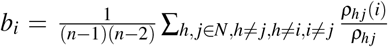, where *ρ*_*h j*_(*i*) is the weights of shortest path between h and j that passes through i.

### Statistical Analysis

We used different statistical methods as described below, and for the multiple testing we relied on the Bonferroni correction^40,41^. Where applicable, the thresholds were obtained by dividing 0.05 by the number of independent tests. We stated the corrected threshold value and the number of independent tests in the caption of each table in the Results section.

#### Quantifying the Change in CDR and Connectivity Metrics

To determine the longitudinal change in CDR, local and global connectivity metrics, we calculated the absolute difference between the first visit (the baseline visit) and the first visit after 12 months (the follow-up visit). Unless stated otherwise, this is the primary way we used to quantifying the longitudinal change in this analysis.

#### Estimation of Gene Expression from Multiple Probe Sets

Different probe set expression values were present for each gene in the data. To estimate a representative gene expression out of the probe set expression, we conducted a non-parametric Mann-Whitney U test to evaluate whether the expression in AD was different from those of healthy elderlies. For each gene, we selected the probe set expression that has the lowest Mann-Whitney U p-value. In this way, we selected the most differentially expressed probe sets in our data and considered those for the remaining analysis.

#### Spearman’s Rank Correlation Coefficient

To test the statistical significance of pair-wise undirected relationships, we used the Spearman’s rank correlation coefficient (*ρ*). The Spearman coefficient is a non-parametric method which ranks pairs of measurements and assesses their monotonic relationship. We report here the coefficient *ρ* along with the corresponding p-value to evaluate the significance of the relationship. A *ρ* of ±1 indicates a very strong relationship, while *ρ* = 0 means there is no relationship.

#### Quantile Regression

To model the directed relationship between two variables, we used the quantile regression model^42^. This model is used as an alternative to the linear regression model when the linear regression assumptions are not met. This fact allows the response and predictor variables to have a non-symmetric distribution. The quantile regression model estimates the conditional median of the dependent variable given the independent variables. Besides, it can be used to estimate any conditional quantile; and is therefore robust to outliers. Accordingly, in this work we used the quantile regression to model the directed relationship between two variables. As an alternative to the linear regression, we chose the 50^*th*^ percentile (the median) as our quantile and estimated the conditional median (rather than the conditional mean in case of the ordinary linear regression) of the dependent variable across given values of the independent variable.

#### Ridge Regression

For estimating the relationship between more than two variables, we used ridge regression^43^. The basic idea behind this model is that it solves the least square function penalizing it using the *l*_2_ norm regularization. More specifically, the ridge regression minimizes the following objective function: 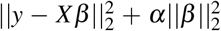,i.e., 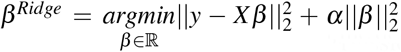 where *y* is the dependent (or response) variable, *X* is the independent variable (feature, or predictor), *β* is the ordinary least square coefficient (or, the slope), *α* is the regularization parameter, *β* ^*Ridge*^ is the ridge regression coefficient, *argmin* is the argument of minimum and it is responsible for making the function attain the minimum and is *L*_2_(*v*) = ||*v*||_2_ which represents the L2 norm function^44^. For example, the CDR scores are the response variables; and brain connectivity features or gene expressions are the predictor variables. We normalized the predictors to get a more robust estimation of our parameters.

#### Software

We used python 3.7.1 for this work; our code has been made available under the MIT License https://choosealicense.com/licenses/mit/, and is accessible at https://github.com/elssam/RGLCG.

## Results

### Descriptive Statistics

Initially, we used descriptive statistics plots to visualize the data for the two populations; the AD and matched control subjects. To facilitate the integrated analysis, we looked into the different sets of data individually to have a better understanding of the underlying statistical distribution, and chose the best analysis methods accordingly. Firstly, we plotted the global and local connectivity metrics in a way that illustrates the longitudinal change between the two visits (i.e. baseline and follow-up visits). The follow-up visit was one year after the baseline screening. The global connectivity metric box plots show the baseline and follow-up distributions for both AD and healthy elderly for transitivity, Louvain modularity, characteristic path length and global efficiency (Figure 1). Figure 1 shows that the global longitudinal changes in connectome metrics are statistically significant among the AD subjects and not mere artifacts, but not within the control population which has non significant changes. In other words, comparing the two groups (AD and healthy elderly) in terms of the change in global connectivity metrics overtime, the only significant differences found between baseline and follow-up were within the AD group. These were found in the characteristic path length (p-value 0.0057), global efficiency (p-value 0.0033), and Louvain modularity (p-value 0.0086). The test used here was the Wilcoxon signed rank test for paired samples with a threshold of 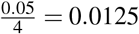, where the division by 4 is due to the multiple hypothesis correction.

**Figure 1.**
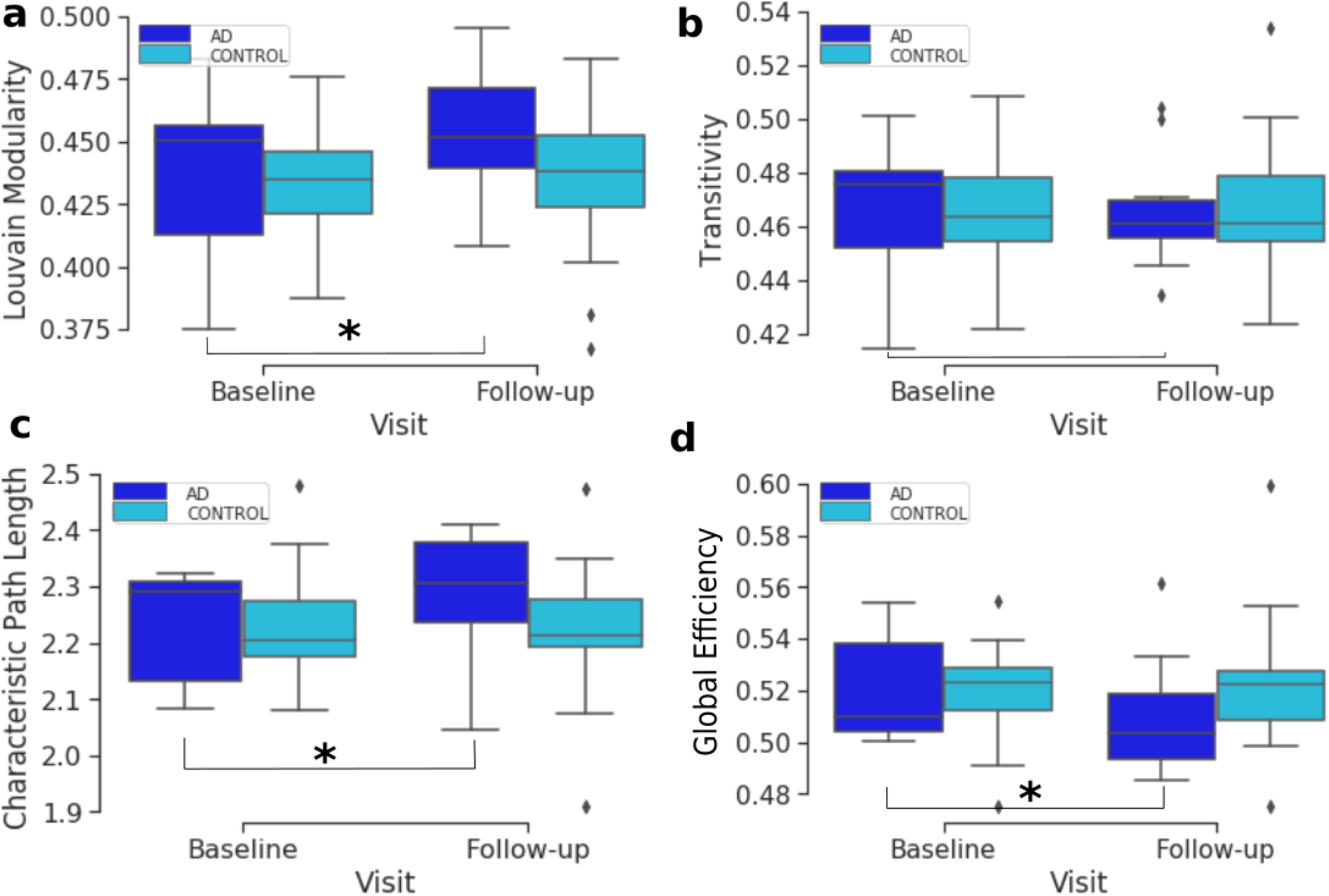
Box plots showing the distribution of brain segregation and integration global connectivity metrics. The plots compare the baseline and follow-up distributions for AD and healthy elderly subjects for Louvain modularity (a), transitivity (b), characteristic path length (c) and global efficiency (d). The asterisk denotes a significant change between the baseline to the follow-up visit.

Supplementary Figures **??, ??** and **??** show the distribution of the absolute differences between the two visits in local efficiency, clustering coefficient and betweenness centrality connectivity metrics, respectively, at each of the AAL brain regions. These figures show that the amount of change in local connectivity metrics varies across the AAL regions. A list of the brain atlas region names and abbreviations is available in Supplementary Table **??**. Moreover, in Supplementary Figure **??**, we show the scatter and violin plots of the six CDR scores, at the baseline and follow-up. The CDR scores are explained in the Materials and Methods section. As such, both global and local connectivity metrics show non-symmetric distribution in the baseline, follow-up and absolute change between them (see Figure 1, Supplementary Figures **??, ??** and **??**). Therefore, we will use non-parametric models and statistical tests in the following analysis.

Comparing the three local connectivity features to each other using Spearman correlation, as shown in Figure 3, we observe a high redundancy between local efficiency and cluster coefficients, assuming they are computed at the same time point. Moreover, we analysed the relationship between the above-mentioned connectivity metrics and other related measures (e.g degree centrality). As shown in Figure 2, the correlation between nodal degree and betweenness centrality appears to be low, whereas there is a negligible correlation between degree centrality and both local efficiency and clustering coefficient.

**Figure 2.**
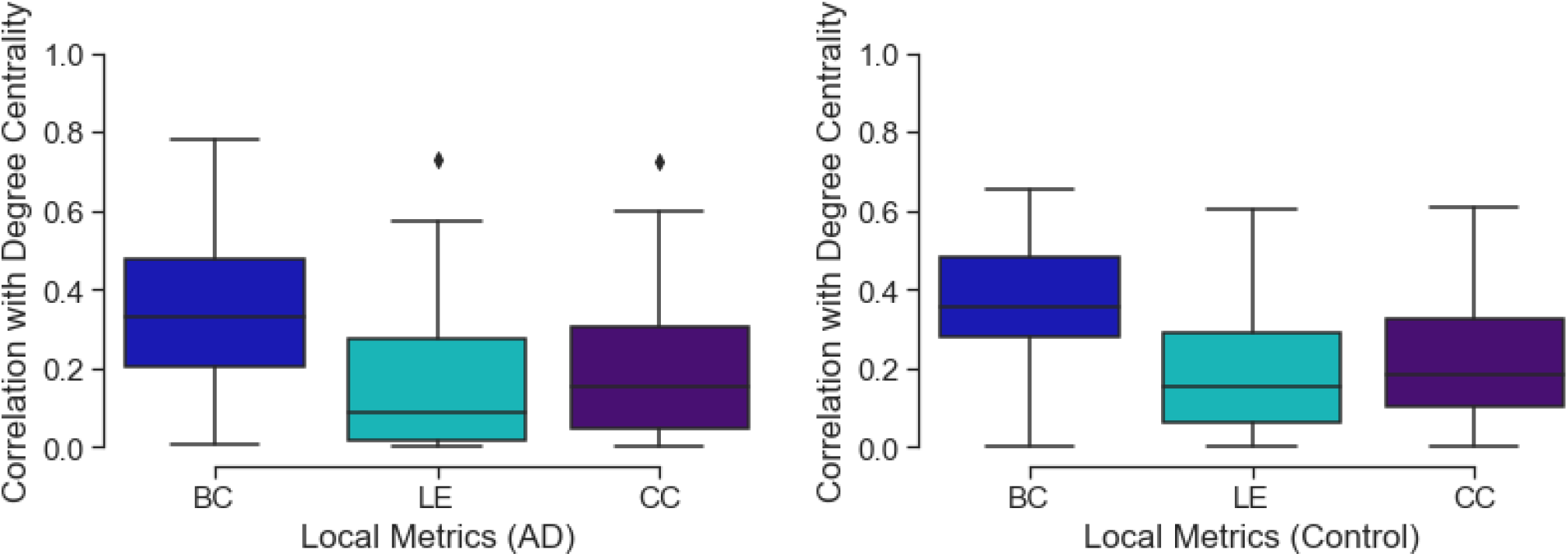
The figure shows three box plots representing the distribution of correlation between degree centrality and the three local connectivity metrics among AD subjects (left) control (right). Each boxplot represents the correlation values between the degree centrality of a node and the related local efficiency (LE), betweenness centrality (BC) and clustering coefficient (CC). Values close to 0 represent low correlation while those close to 1 show a high correlation.

**Figure 3.**
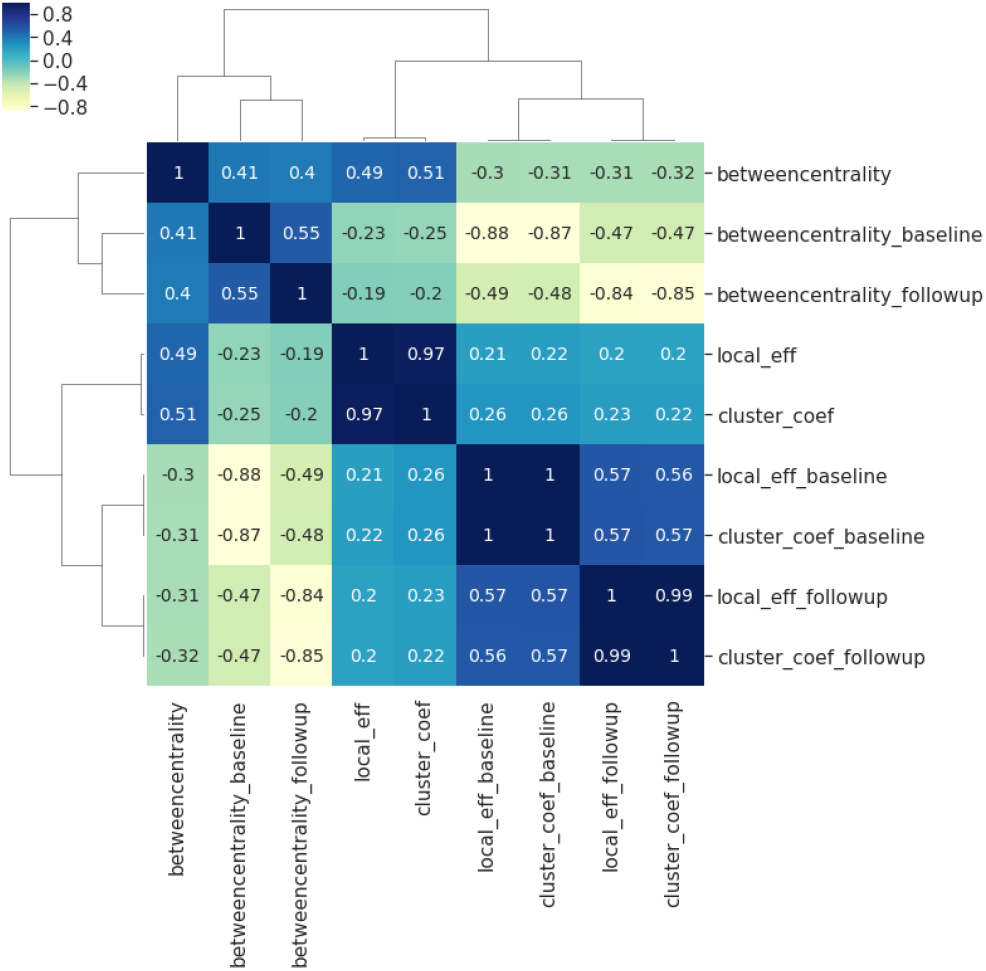
Spearman correlations between the three local connectivity metrics; local efficiency, clustering coefficient and betweenness centrality, at baseline (suffix: _baseline), follow-up (suffix:_followup) and the absolute difference between them (no suffix). The calculation of Spearman’s coefficient combines all 90 brain regions for the combined sample (i.e. both AD and control). The plot shows a very strong relationship between clustering coefficient and local efficiency at baseline, follow-up and the absolute difference between the two visits.

### Gene Expression

We derived a list of 17 AD risk factor genes from BioMart, and retrieved 56 related probes sets from ADNI data. We performed a Mann-Whitney U test which aims at testing whether a specific probe set’s expression is different between AD and healthy controls. For each gene, we chose the probe set that has the lowest p-value. Table 1 reports the selected probe set (with the minimum p-value) at each gene. It is worth mentioning that we are using probe sets, rather than single probes. These probe sets represent different parts of the transcripts rather than representing single alternative splicing of the gene, and are normalised across probes for each gene. Expression values were normalised using the robust multi-chip average method in the Affymetrix U219 array, which consists of a total number of 49,293 probe sets and 530,467 probes, as explained in the Genetic Data Acquisition section.

**Table 1.** Mann-Whitney U test top results for the difference between AD and controls in probe set expression

| Gene | Top results |  |  |
| --- | --- | --- | --- |
|  | Chromosome | Probe set id | p-value |
| *APBB2* | 4 | 11734823_a_at | 0.02575 |
| *MPO* | 17 | 11727442_at | 0.38631 |
| *APP* | 21 | 11762804_x_at | 0.01396 |
| *ACE* | 17 | 11752871_a_at | 0.24478 |
| *PLAU* | 10 | 11717154_a_at | 0.01396 |
| *PAXIP1* | 7 | 11755176_a_at | 0.45499 |
| *HFE* | 6 | 11736346_a_at | 0.11881 |
| *SORL1* | 11 | 11743129_at | 0.10912 |
| *A2M* | 12 | 11715363_a_at | 0.28592 |
| *NOS3* | 7 | 11725467_a_at | 0.04261 |
| *BLMH* | 17 | 11757556_s_at | 0.09356 |
| *ADAM10* | 15 | 11751180_a_at | 0.14278 |
| *PLD3* | 19 | 11715382_x_at | 0.17304 |
| *ApoE* | 19 | 11744068_x_at | 0.05962 |
| *PSEN1* | 14 | 11718678_a_at | 0.29453 |
| *PSEN2* | 1 | 11723674_x_at | 0.04862 |
| *ABCA7* | 19 | 11755091_a_at | 0.45499 |

After estimating the expression of the 17 genes, as explained in the Materials and Methods, we plotted a heatmap of the gene expression profiles shown in Supplementary Figure **??**. Some of the genes appear to be highly expressed (e.g. *SORL1* and *PSEN1*), while others show very low expression (e.g. *HFE* and *ACE*). To avoid double-dipping in estimating the effect size of gene expression, the subsequent analysis will not depend on disease status (i.e. AD and control), but rather, on quantitative measurements (e.g. local and global connectivity metrics) derived from the whole sample.

### Association Analysis

We studied the undirected associations of the 17 gene expression values with the longitudinal change in global and local brain connectivity, as well as the associations with longitudinal CDR and connectivity changes. Firstly, we performed an association analysis of gene expression with the connectivity changes locally, at each AAL brain region. In Table 2 we show the top results reported along with the Spearman correlation co-efficient. The *APP* gene (*ρ* =-0.58, p-value=1.9e-05) and *BLMH* (*ρ* =0.57, p-value=2.8e-05) are the top genes in the list, and although they both did not hit the significance threshold, they show potential association with the change in local efficiency at the right middle temporal gyrus (Temporal_Mid_R AAL region) and clustering coefficient at the left Heschl gyrus (Heschl_L), respectively. Supplementary Figure **??** shows the scatter plots related to these significant associations.

**Table 2.** Top results of Spearman associations between AD gene expression and local connectivity metrics.

| Gene | Sorted by P-value. Threshold = $\frac{0.05}{17 \times 90 \times 3} = 1.09e-05$ | | | | |
| --- | --- | --- | --- | --- | --- |
| | Region | Region id | Metric | $\rho$ | P-value |
| *APP* | Temporal_Mid_R | 86 | local_eff | -0.5805 | 1.9e-05 |
| *BLMH* | Heschl_L | 79 | cluster_coef | 0.5708 | 2.8e-05 |
| *PSEN1* | Occipital_Mid_R | 52 | b_centrality | -0.5598 | 4.3e-05 |
| *BLMH* | Heschl_L | 79 | local_eff | 0.5591 | 4.4e-05 |
| *APP* | Temporal_Mid_R | 86 | cluster_coef | -0.5197 | 0.000182 |
| *PAXIP1* | Amygdala_L | 41 | cluster_coef | 0.5064 | 0.000281 |
| *PLAU* | Angular_R | 66 | cluster_coef | 0.484 | 0.000567 |
| *PLAU* | Angular_R | 66 | local_eff | 0.4838 | 0.00057 |
| *ACE* | Postcentral_L | 57 | b_centrality | 0.4648 | 0.000998 |
| *ADAM10* | Postcentral_L | 57 | local_eff | -0.4602 | 0.001133 |
| *PAXIP1* | Parietal_Sup_L | 59 | b_centrality | -0.4585 | 0.00119 |
| *PLAU* | Fusiform_L | 55 | b_centrality | 0.4564 | 0.001262 |
| *SORL1* | Putamen_R | 74 | local_eff | -0.4528 | 0.001395 |
| *PSEN2* | Frontal_Inf_Oper_R | 12 | local_eff | 0.4457 | 0.001693 |
| *PLAU* | Frontal_Inf_Oper_L | 11 | b_centrality | 0.4454 | 0.001704 |
| *ABCA7* | Temporal_Inf_L | 89 | local_eff | 0.442 | 0.001866 |

In Table 2, there is a similar pattern observed in association results between the regional clustering coefficient and local efficiency, e.g. both metrics are associated with *BLMH* at the left Heschl gyrus (Heschl_L), *APP* at the right middle temporal gyrus (Temporal_Mid_R) and *PLAU* at the right angular gyrus (Angular_R). We interpret this by the strong correlation that exists between the local efficiency and clustering coefficient at the baseline, follow-up and also, the absolute change (see Figure 3). On the other hand, Table 3 reports the top results of the association between gene expression and the change in brain global connectivity. In this case, all observed associations were not statistically significant after correcting for multiple hypotheses (threshold is stated in Table 3).

**Table 3.** Top 20 Spearman association results of the change in global network metrics with targeted AD gene expressions.

| Gene | Threshold = $\frac{0.05}{17 \times 4} = 7.35e - 04$ | | |
| --- | --- | --- | --- |
| | $\rho$ | P-value | Global Feature |
| *PAXIP1* | 0.3889 | 0.0069 | transitivity |
| *PLAU* | -0.3824 | 0.008 | global_eff |
| *ACE* | -0.3696 | 0.0106 | transitivity |
| *PLAU* | -0.3523 | 0.0151 | char_path_len |
| *ABCA7* | -0.3492 | 0.0161 | char_path_len |
| *PSEN1* | -0.299 | 0.0412 | transitivity |
| *APP* | -0.2602 | 0.0774 | char_path_len |
| *PLAU* | -0.2542 | 0.0847 | Louvain |
| *ApoE* | -0.2506 | 0.0893 | char_path_len |
| *ADAM10* | -0.2365 | 0.1095 | Louvain |
| *ACE* | 0.2291 | 0.1213 | Louvain |
| *NOS3* | -0.2207 | 0.1359 | char_path_len |
| *NOS3* | -0.2202 | 0.1369 | global_eff |
| *ABCA7* | -0.2164 | 0.1441 | global_eff |
| *HFE* | -0.2012 | 0.1752 | char_path_len |

### Regression Analysis: the Change in Local and Global Brain Connectivity with Gene Expression

We analyzed the directed association through regressing the change in local connectivity (as a dependant variable), at each AAL region, on gene expression using (as an independent variable or predictor) a quantile regression model. Table 4 reports the top results, along with the regression coefficient, p-values and t-test statistic. *PLAU* was the only significant gene, affecting the absolute change in betweenness centrality at left Fusiform gyrus (Fusiform_L) with an increase of 487.13 at each unit increase in *PLAU* expression (p-value= 3*e −* 06).

**Table 4.**
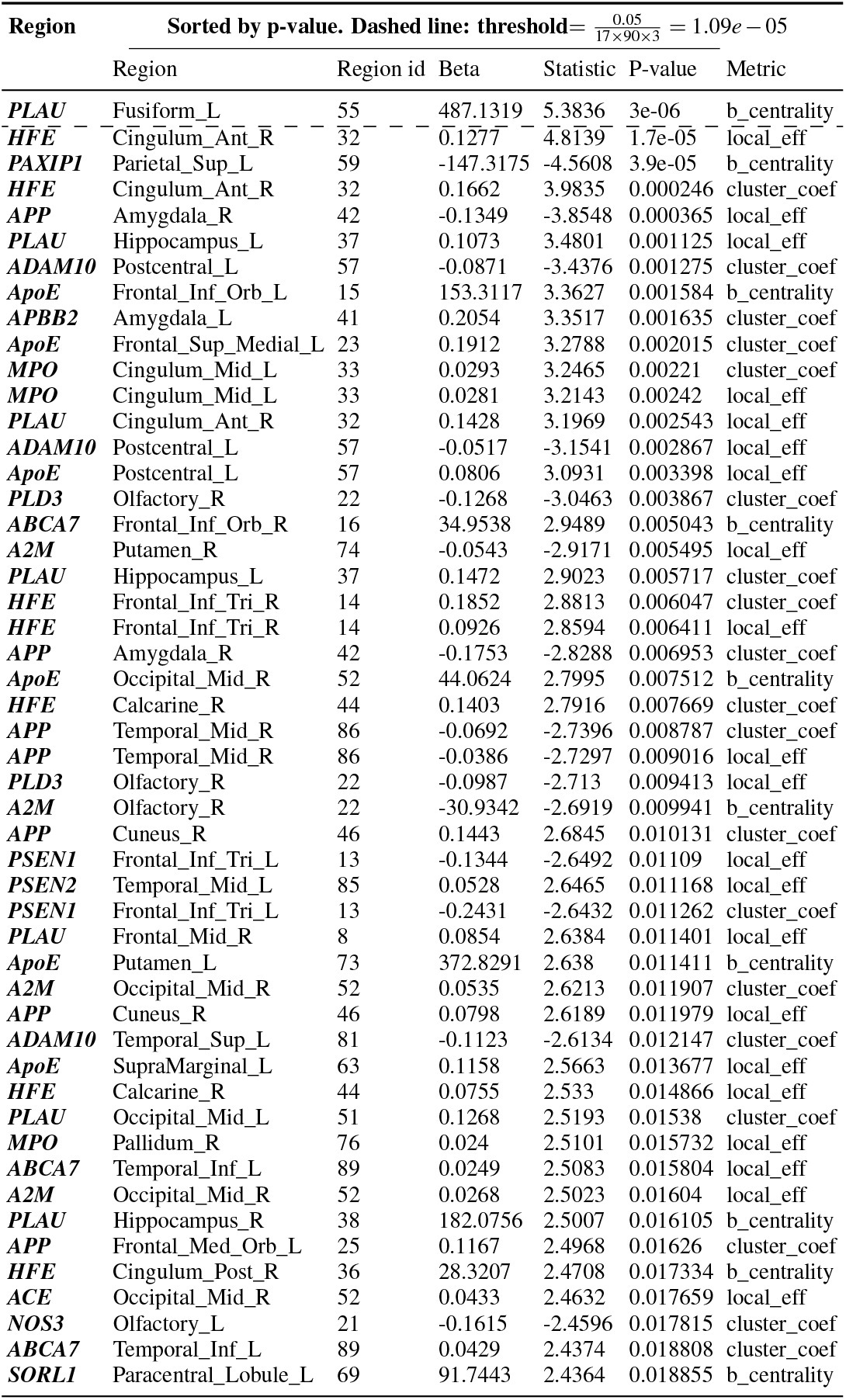
Top 50 quantile regression results of the change in local network metrics (*y*) and targeted AD gene expression (*x*)

This was followed by the expression of *HFE* with an effect size of 0.1277 on the change in local efficiency at the right anterior cingulate and paracingulate gyri (Cingulum_Ant_R). Those observed associations are illustrated in Figure 4, and, the protein-protein interaction^45^ of the aforementioned genes are shown in Figure 5. Supplementary Figures **??, ??** and **??** show the Manhattan plots for the -log10 of the p-values corresponding to the quantile regression models of the change in local efficiency, clustering coefficient and betweenness centrality, respectively.

**Figure 4.**
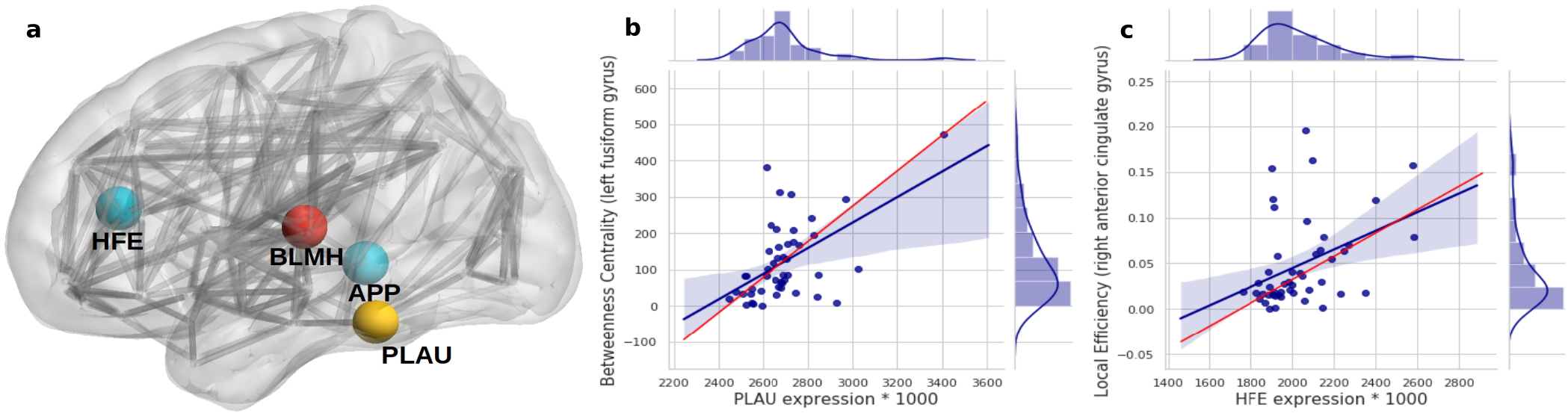
Subfigure (a) higlights regions in the brain where significant (and potential) associations between gene expression and longitudinal change in local connectivity metrics were found, using quantile regression (*HFE* and *PLAU*) and Spearman associations (*APP* and *BLMH*). Each gene is plotted at the AAL brain region where the association was significant (or nearly significant); *APP* at Temporal_Mid_R, *BLMH* at Heschl_L, *PLAU* at Fusiform_L and *HFE* at Cingulum_Ant_R. (b) and (c) are scatter plots to visualize the association between *PLAU* gene expression and betweenness centrality in the left fusiform gyrus (a), and between the expression of *HFE* gene with local efficiency in right anterior cingulate gyrus (b). The red line on the plots represents the median (quantile) regression line, while the blue line represents the ordinary least square line. It is important to bear in mind that after the multiple hypothesis testing only the PLAU gene was still significant.

**Figure 5.**
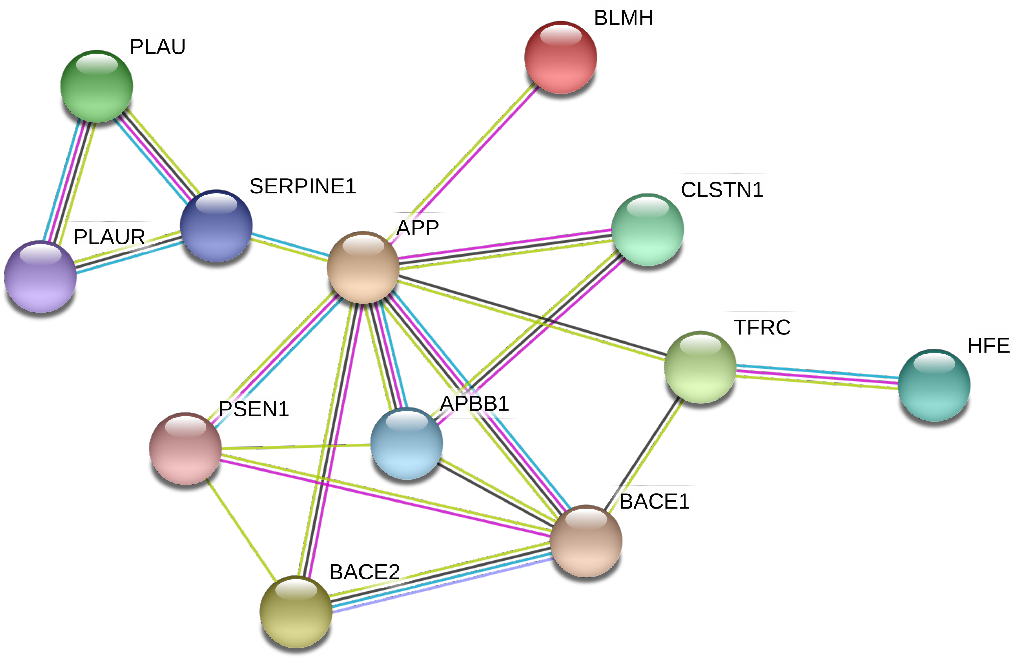
Protein-protein interactions between the genes with significant (or potential) correlation to brain connectivity metrics. It is evident that the genes interact either directly with APP or via one intermediate node/gene. In this sub-network extracted from STRING^45^, the different color lines represents different types of interaction: Cyan edges are interactions from curated databases, purple are experimentally determined, yellow are from text mining, and black are from co-expression data.

Similarly, we regressed the absolute change of global connectivity measures on gene expression values and the top results are shown in Table 5.

**Table 5.** Top quantile regression results of the change in global network metrics and targeted AD gene expressions.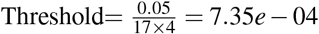.

| Gene | Threshold $\frac{0.05}{17 \times 4} = 7.35e-04$ | | | |
| --- | --- | --- | --- | --- |
|  | Beta | Statistic | P-value | Metric |
| *PAXIP1* | 0.0155 | 2.1179 | 0.039738 | transitivity |
| *PSEN1* | -0.0366 | -2.0821 | 0.04305 | transitivity |
| *A2M* | 0.0178 | 1.9728 | 0.054679 | Louvain |
| *PLAU* | -0.0157 | -1.9288 | 0.06008 | global_eff |
| *APBB2* | 0.0166 | 1.7579 | 0.085573 | Louvain |
| *ABCA7* | 0.0077 | 1.5185 | 0.135881 | transitivity |
| *BLMH* | 0.0101 | 1.3476 | 0.184529 | transitivity |
| *ACE* | -0.0162 | -1.2618 | 0.213524 | transitivity |
| *ADAM10* | -0.0147 | -1.2538 | 0.21638 | Louvain |
| *ApoE* | -0.0555 | -1.1673 | 0.249246 | char_path_len |
| *PLD3* | -0.0096 | -1.1433 | 0.258978 | transitivity |
| *ABCA7* | -0.0156 | -1.1367 | 0.261684 | char_path_len |
| *SORL1* | 0.0147 | 1.1278 | 0.265372 | transitivity |
| *ApoE* | -0.0099 | -1.1048 | 0.275101 | global_eff |
| *HFE* | -0.0161 | -1.0436 | 0.302239 | Louvain |

### Additive Genetic Effect on Brain Regions

To visualize the overall contribution of AD gene risk factors used in this work on distinct brain areas, we added up the -log10 p-values for the gene expression coefficients at each of the 90 AAL regions. The p-values were obtained from the quantile regression analysis between the gene expression values and each of the three connectivity metrics - those are the absolute difference between baseline and follow-up of local efficiency, clustering coefficient and betweenness. Figure 6 summarizes this by 1) representing the brain connectome without edges for each of the connectivity metrics, 2) each node represents a distinct AAL region and is annotated with the name of the region, 3) the size of each node is the sum-log10 of the regression coefficient associated p-vales for all the genes. It is clear from Figure 6 that local efficiency and clustering coefficient show more similar patterns of association with genes compared to betweenness centrality. This means that the gene expression has stronger association with local structure of the brain when using clustering measures (e.g. clustering coefficients), while the pattern of association tends to be weaker when using measures expressing the state of a region being between two others (e.g. betweenness centrality).

**Figure 6.**
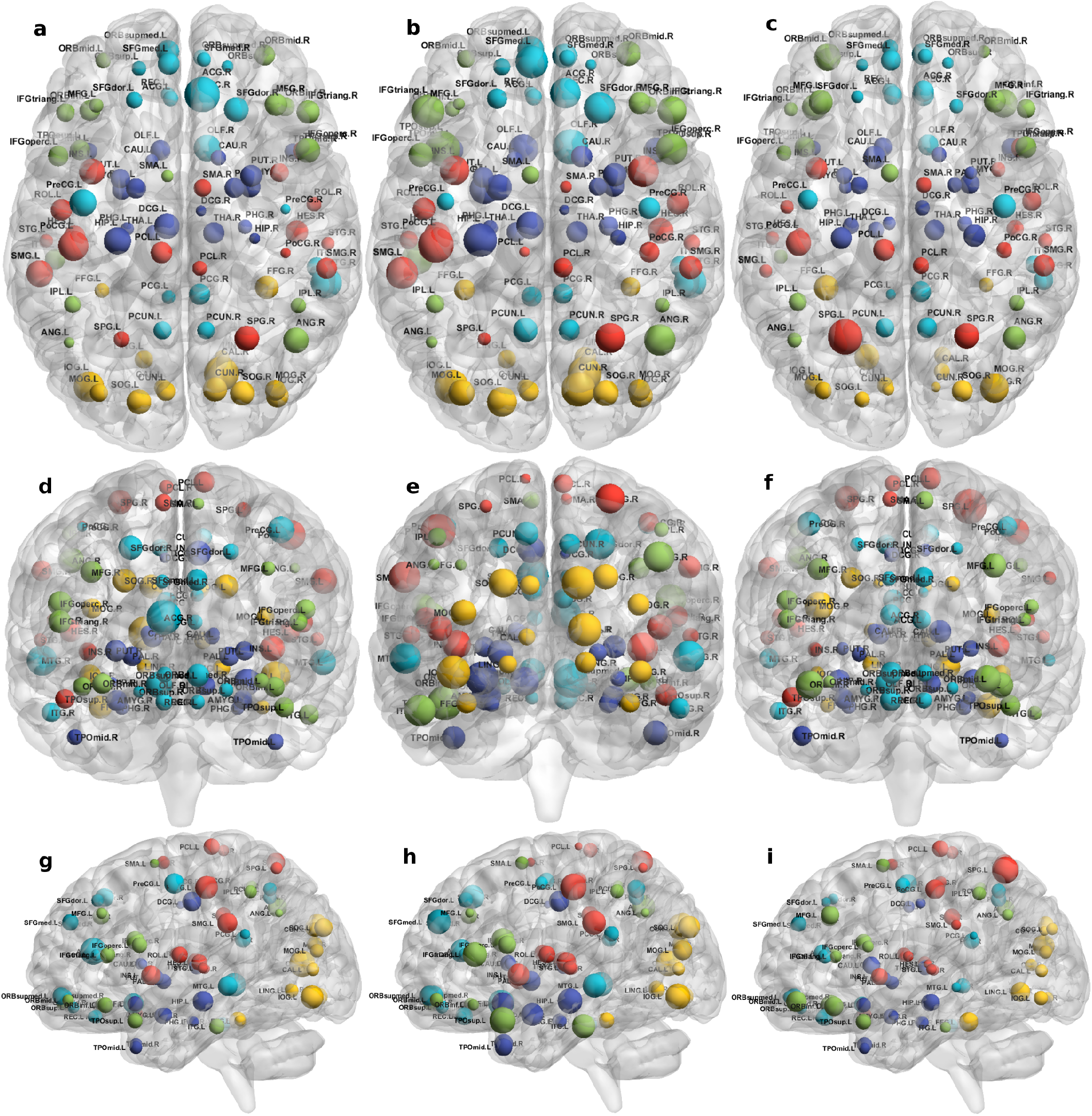
Connectome representations showing the local connectivity additive genetic effect at each AAL node. The subfigures show the axial (top; (a), (b) and (c)), coronal (middle; (d), (e) and (f)), and sagittal (bottom; (g), (h) and (i)) planes of the brain, the node size represents the local efficiency (left; (a), (d) and (g)), clustering coefficient (middle; (b), (e) and (h)) and betweenness centrality (right; (c), (f) and (i)). Colors of the nodes are automatically assigned by the BrainNet Viewer. The acronyms of the brain regions are explained in Supplementary Table **??**.

The colors are assigned automatically by the BrainNet Viewer. Overall, although the gene contributions to the absolute change in local efficiency have a similar pattern to that of clustering coefficient, the contribution to betweenness centrality change is relatively small.

### Regression Analysis: the difference in CDR with the difference in Global and Local Connectivity

To asses the directed and undirected association of the longitudinal measures of global connectivity and CDR scores, we calculated the difference between baseline and follow-up visits for both CDR and global connectivity metrics, i.e. *CDR*_*baseline*_ *− CDR*_*follow−up*_ and *metric*_*baseline*_ *− metric*_*follow−up*_, respectively. The Spearman and quantile regression results are both shown in Table 6. We observed a correlation between the increase of the transitivity score (global brain segregation) and the CDR memory score over time (*β* = *—* 6.14*e−*06, p-value= 0.0034). On the other hand, there is a positive correlation between global efficiency (global brain integration) and the CDR “home and hobbies” score.

**Table 6.** Quantile regression results of the difference in CDR (y) with the difference in global connectivity (x)

| Global Metric | Threshold=$\frac{0.05}{6 \times 4} = 2.08e - 3$. * represents significant p-value. | | | | | |
| --- | --- | --- | --- | --- | --- | --- |
|  | CDMEMORY | CDORIENT | CDJUDGE | CDCOMMUN | CDHOME | CDCARE |
| **Q. Regression:** | $\beta$ (p-value) | | | | | |
| transitivity | -6.14e-06 (0.0034*) | -1.8e-06 (nan) | -3.2e-07 (nan) | 8.4e-07 (0.9249) | 4.13e-06 (0.7324) | -3.1e-07 (nan) |
| global_eff | 1.3e-06 (0.8572) | 3.36e-06 (0.9944) | 3.5e-07 (0.9826) | 3.09e-06 (0.3131) | 9.5e-06 (0.1613) | -2.5e-07 (nan) |
| Louvain | -2.64e-06 (0.1683) | -6.84e-06 (0.0352) | 1.21e-06 (0.0012) | -1.01e-06 (0.8125) | -2.16e-06 (0.9424) | -6.3e-07 (0.1787) |
| char_path_len | -5.8e-07 (0.8562) | -8.3e-07 (0.8791) | -8e-08 (0.8637) | -8.3e-07 (0.0361) | -2.14e-06 (0.0442) | -2e-08 (nan) |
| **Spearman:** | $\rho$ (p-value) | | | | | |
| transitivity | -0.3395 (0.021) | -0.0763 (0.6142) | -0.0618 (0.6835) | 0.0661 (0.6623) | 0.161 (0.2851) | -0.0081 (0.9574) |
| global_eff | 0.0483 (0.7497) | 0.0056 (0.9707) | 0.0505 (0.7388) | 0.2685 (0.0712) | 0.4145 (0.0042*) | -0.0263 (0.8625) |
| Louvain | -0.1955 (0.1928) | -0.3119 (0.0349) | 0.2077 (0.166) | -0.0968 (0.522) | -0.11 (0.4666) | -0.0628 (0.6784) |
| char_path_len | -0.1183 (0.4337) | -0.0516 (0.7333) | -0.0281 (0.8531) | -0.2909 (0.0498) | -0.3811 (0.009) | -0.0065 (0.9658) |

Similarly, in Supplementary Table **??** we looked at the monotonic effect of local connectivity metrics on the seven CDR scores, both represented as the subtraction of the follow-up visit from the baseline visit. The increase in betweenness centrality was shown to have different effects on the CDR score over the one-year time period. For example, as the betweenness centrality decreases over time, the judgement and problem solving increases in severity by 1.06e-08 over time (p-value=1.32e-17), in the frontal lobe (Frontal_Inf_Oper_L).

### Multivariate Analysis: Ridge Regression

We regressed the difference in CDR visits (response variable; Y), one score at a time, on both the difference in global brain connectivity (predictor; X1), one connectivity metric at a time and all gene expression values (predictor; X2), using the ridge regression model. Supplementary Table **??** reports the mean squared error (the score column) and shows the top hits in the multiple ridge regression. It shows that the *α* (alpha column) could not converge, using the cross-validation, when the response variables were judgment or personal care. However, the CDR score results show that genes and connectivity metrics have a small effect (*β*) on the response variables (the change in CDR scores over time), and the larger effects were observed when using the total CDR score (CDR_diff) as a response variable. The connectivity metrics and expression of genes have both negative and positive effects on CDR change. The expression of *ApoE*, for example, has a negative effect (*β*) of *−* 0.24 on the change in memory score, i.e. the memory rating decreases by 0.24 as the *ApoE* expression increases. While when the expression of *ApoE* increases one unit, the home and hobbies score increases, over time, by 0.12.

## Discussion

In this paper we aimed to analyse the longitudinal change of local and global brain connectivity metrics in Alzheimer’s Disease, using a dataset from ADNI. We evaluated the association between the most commonly used connectivity metrics and the expression of known AD risk genes. Finally, we studied the multivariate association between connectivity metrics, gene expression and dementia clinical ratings. It is worth mentioning that it is not practical to do expression analysis on brain tissues, thus, we had to use a proxy for what is generally happening in the body by using the blood samples. This is the closest proxy that is feasible when working with both patients and controls, also in the view of using the proposed biomarkers in daily practice. Moreover, we verified that the found genes were expressed in the brain parenchyma by using the the ISH Allen Atlas^46^, which confirmed positively their expression.

Our results show that Alzheimer’s risk genes can manipulate the amount of change observed in the structural connectome, measured as the absolute difference of longitudinal connectivity metrics. We also show that longitudinal regional connectivity metrics, global brain segregation and integration have effects on the CDR scores. More specifically, as the disease progresses, we observe a correlation between brain segregation and cognitive decline, the latter is measured as the memory CDR score. We also noticed a correlation between the global efficiency and the home and hobbies CDR scores (see Table 6). Furthermore, we observe a consistent decrease, over time (though did not hit the significance threshold), in the local efficiency (a network connectivity metric of a node is defined as the inverse of the shortest average path between any pair of that node’s neighbors^6^) at the right middle temporal gyrus (Temporal_Mid_R; see Table 2) in response to an increase in *APP* expression. The same connectivity metric showed another potential increase over time at the right anterior cingulate and paracingulate gyri (see Table 4) as the expression of *HFE* increases.

Prescott et al.^47^ have investigated the differences in the structural connectome at the three clinical stages of AD using a cross-sectional study design. They targeted regional brain areas that are known to have increased amyloid plaque. Their work suggested that connectome damage might occur at an earlier preclinical stage towards developing AD. Here, we further adapted a longitudinal study design and incorporated known AD risk genes in analysing the changes in the connectome. We specifically focus on how the damage in the connectome is associated with gene expression and how is that change in connectome affects dementia symptoms, globally and locally at distinct brain regions. Aside from our previous work which examined the *ApoE* associations with longitudinal global connectivity in AD using longitudinal global connectivity metrics^8^, this study, to our knowledge, is the first of its type to include gene expression data in a longitudinal analysis of global and local brain connectivity. However, similar work has been done in schizophrenia where the the association between longitudinal magnetic resonance imaging features, derived from the DWI, and higher genetic risk for schizophrenia were investigated using structural brain connectivity^48^.

The expression in the regional areas of the brain where we found significant (i.e. PLAU) or potential (i.e. HFE, APP and BLMH) association with local connectivity (highlighted in Figure 4) are mostly in line with previous findings of the regional molecular properties in the brain (e.g.^49^). In this study we found that the Plasminogen activator, urokinase (*PLAU*) expression affects the betweenness centrality (a measure of the node’s contribution to the flow of information in a network^6^) in the left fusiform gyrus, over time (see Table 4 and Figure 4). Although the functionality of this region is not fully understood, its relationship with cognition and semantic memory was previously reported^50^. *PLAU*, on the other hand, was shown to be a risk factor in the development of late-onset AD as a result of its involvement in the conversion of plasminogen to plasmin - a contributor to the processing of APP by the urokinase-type plasminogen activator^51^.

Among further results - including those not surviving the multiple hypothesis testing - potentially align with findings in the literature of genetics and neuroimaging. Robson et al.^52^ studied the interaction of the *C282Y* allele *HFE* - the common basis of hemochromatosis - and found that carriers of *ApoE*-4, the C2 variant in TF and C282Y are at higher risk of developing AD. The *HFE* gene is also known for regulating iron absorption, which results in recessive genetic disorders, such as hereditary haemochromatosis also related to AD^53^. These studies align with our findings where we show that *HFE* expression (though it did not service the significance threshold after multiple testing correction) can potentially affect the local efficiency at the right anterior cingulate gyrus (see Table 4 and Figure 4), which might indicate a possible effect of *HFE* expression on the personality of AD patients or those at risk of developing the disease. When examining the linear associations between gene expression and local connectivity (see Table 2 and Supplementary Figure **??**), we found that the local efficiency in the right middle temporal gyrus (nearly hit the significance threshold), known for its involvement in cognitive processes including comprehension of language, negatively associates with *APP* expression. We also found a potential correlation between the left Heschl gyrus’s clustering coefficient and the bleomycin hydrolase (BLMH) expression. In the human brain, the BLMH protein is expressed in the neocortical neurons and associated to misfolded proteins related to AD. Some studies^54,55^ have found that a variant in the *BLMH* gene, which leads to the Ile443→ Val in the BLMH protein, increases the risk of AD; this was strongly marked in *ApoE*-4 carriers. The BLMH protein can process the *Aβ* protein precursor, and is involved in the production of *Aβ* peptide^56^.

The regional expression of the *APP* gene has been shown to be positively correlated with the severity of regional amyloid deposition observed in PET studies. The temporal medial region and the fusiform gyrus of the brain are two of the most affected regions by amyloid deposition and have high levels of *APP* expression^49^. Recent findings from tau-sensitive positron emission tomography data have also confirmed the spatial correspondence between accumulation of tau pathology and neurodegeneration in AD patients, within the same regions, though only correlations with the *ApoE* genes were investigated^57^. The BLMH protein alters the processing of APP and significantly increases the release of its proteolytic fragment. It has been previously reported to be expressed and have an impact on the hippocampal tissues, but not investigated in other brain regions^58^. To our knowledge, apart from the general expression in the brain parenchyma reported in the Allen brain atlas^59^, no study has shown spatial expression of the *PLAU* and *HFE* genes among AD patients. Nevertheless, we hypothesize that there is a potential correlation of these gene expression in specific nodes of the brain connectome, and that it is related to their interaction with the *APP* gene which is particularly expressed in the nodes highlighted in Figure 4. This hypothesis is also supported by the protein-protein interaction shown in Figure 5.

Even though none of the AD risk genes showed a significant effect on the longitudinal change in global connectivity (see Tables 3 and 5), some of these genes showed significant effects on local connectivity changes at specific brain areas (see Table 4 and Table 2). The global connectivity metrics of the brain, on the other hand, have shown promising results in affecting the change observed in CDR scores, including memory, judgement and problem solving, as well as home and hobbies, as shown in Table 6. Previous work studied the association between genome-wide variants and global connectivity in AD, represented as brain integration and segregation, and found that some genes affect the amount of change observed in global connectivity measures^8^. This suggests that a generalisation of the current study at a gene-wide level might warrant further analysis.

Considering the possible redundancy of brain connectivity metrics^27^ studied here, we looked for correlations with the nodal degrees and other features, and observed a relative correlation with betweenness centrality (see Figure 2). When we compared all the metrics to each other, we only found a correlation between local efficiency and cluster coefficient metrics. Therefore, we hypothesize that the degree is a simpler representation than betweenness centrality that could be used as a substitute. However, this was not the case with other metrics. We suggest either clustering coefficient or local efficiency should also be investigated in similar studies. More generally, when correcting our threshold, we did not consider the number of metrics, because of the high co-linearity. Therefore, we believe that reporting all, or only one, did not affect the results.

Our work provides new insights into the progression of Alzheimer’s Disease, though replication on a larger sample size is required. Indeed, one limitation here was the small sample size available, as we needed to narrow down our selection of participants to those who 1) attended both the baseline and follow-up visits, 2) have their CDR scores measured, 3) have their genetic and imaging information available. Another limitation is the use of only two time points, the baseline and the first follow-up visits. Other datasets do not offer all those data, and indeed even recent studies have been published with similar sample sizes^60^. Moreover, the limited availability of samples does not allow capturing the effects of connectivity changes over a longer-term or studying the survival probabilities in AD. Extending to more time points would have been useful, but it would have reduced the dataset further. We foresee future work in using a more complex unified multi-scale model to facilitate studying the multivariate effect of clinical and genetic factors on brain diseases, besides considering the complex interplay of genetic factors^61^. Nevertheless, previous studies with similar sample sizes have been able to provide relevant insights for the gene and brain interaction networks^29^. This analysis was conducted in AD, using a longitudinal study design, and highlighted the role of *PLAU, HFE, APP* and *BLMH* in affecting the pattern information propagated in particular regions of the brain, which might have a direct effect on a person’s recognition and cognitive abilities. The four genes were previously shown to be expressed in both the temporal and visual cortex in AD, according to the Allen Human Brain Atlas https://human.brain-map.org/ish/search. Furthermore, the results illustrated the effect of brain structural connections on memory and cognitive process, especially the capacity for reaching a decision or drawing conclusions. The findings presented here might provide better *in-vivo* estimation of local neurodegneration and related treatments. The Braak staging hypothesis is still controversial, and other *in-vivo* studies have shown regional effects using positron emission tomography (PET) imaging (temporal lobe, the anterior cingulate gyrus, and the parietal operculum)^49^. These studies have also shown an impact on the default mode network connectivity^62^. Therefore, further investigation of regional patterns is relevant. Very recent results on treatments in mice showed that drug based modulated neuronal activity can reduce amyloid plaques in specific locations and circuitry^28^. In view of future treatments based on specific spatial location and genetic influences, our study provides some initial insights into connectivity outcomes, and to some extent, enhances our understanding of the regions/circuits that show amyloid aggregation or neurodegeneration.

## Acknowledgements

We would like to acknowledge our funders, the Organisation for Women in Science for the Developing World (OWSD), the Swedish International Development Cooperation Agency (SIDA) and the University of Cape Town for their continuous support. Data collection and sharing for this project was funded by the Alzheimer’s Disease Neuroimaging Initiative (ADNI) (National Institutes of Health Grant U01 AG024904) and DOD ADNI (Department of Defense award number W81XWH-12-2-0012). ADNI is funded by the National Institute on Aging, the National Institute of Biomedical Imaging and Bioengineering, and through generous contributions from the following: AbbVie, Alzheimer’s Association; Alzheimer’s Drug Discovery Foundation; Araclon Biotech; BioClinica, Inc.; Biogen; Bristol-Myers Squibb Company; CereSpir, Inc.; Cogstate; Eisai Inc.; Elan Pharmaceuticals, Inc.; Eli Lilly and Company; EuroImmun; F. Hoffmann-La Roche Ltd and its affiliated company Genentech, Inc.; Fujirebio; GE Healthcare; IXICO Ltd.;Janssen Alzheimer Immunotherapy Research & Development, LLC.; Johnson & Johnson Pharmaceutical Research & Development LLC.; Lumosity; Lundbeck; Merck & Co., Inc.;Meso Scale Diagnostics, LLC.; NeuroRx Research; Neurotrack Technologies; Novartis Pharmaceuticals Corporation; Pfizer Inc.; Piramal Imaging; Servier; Takeda Pharmaceutical Company; and Transition Therapeutics. The Canadian Institutes of Health Research is providing funds to support ADNI clinical sites in Canada. Private sector contributions are facilitated by the Foundation for the National Institutes of Health (www.fnih.org). The grantee organization is the Northern California Institute for Research and Education, and the study is coordinated by the Alzheimer’s Therapeutic Research Institute at the University of Southern California. ADNI data are disseminated by the Laboratory for Neuro Imaging at the University of Southern California.

## Author Contributions

S.S.M.E. conducted the integrated imaging-genetics analysis and wrote the paper. A.C. preprocessed all neuroimaging data and set the connectome pipeline. N.J.M. and A.C. gave constructive feedback on this work continuously and followed up the analysis and writing progress closely. E.R.C. gave feedback on the overall paper. The final version of the paper was proofread by all authors.

## Additional Information

### Supplementary information

Supplementary materials available for this paper.

### Data Availability

The data used in this work are available at the ADNI repository (http://adni.loni.usc.edu/).

