## Supplementary Materials for "Relating Global and Local Connectome Changes to Dementia and Targeted Gene Expression in Alzheimer’s Disease"

### Supplementary Materials for: Relating Global and Local Connectome Changes to Dementia and Targeted Gene Expressions in Alzheimer's Disease

**Samar S. M. Elsheikh, Emile R. Chimusa, Alzheimer's Disease Neuroimaging Initiative, Nicola J. Mulder and Alessandro Crimi**

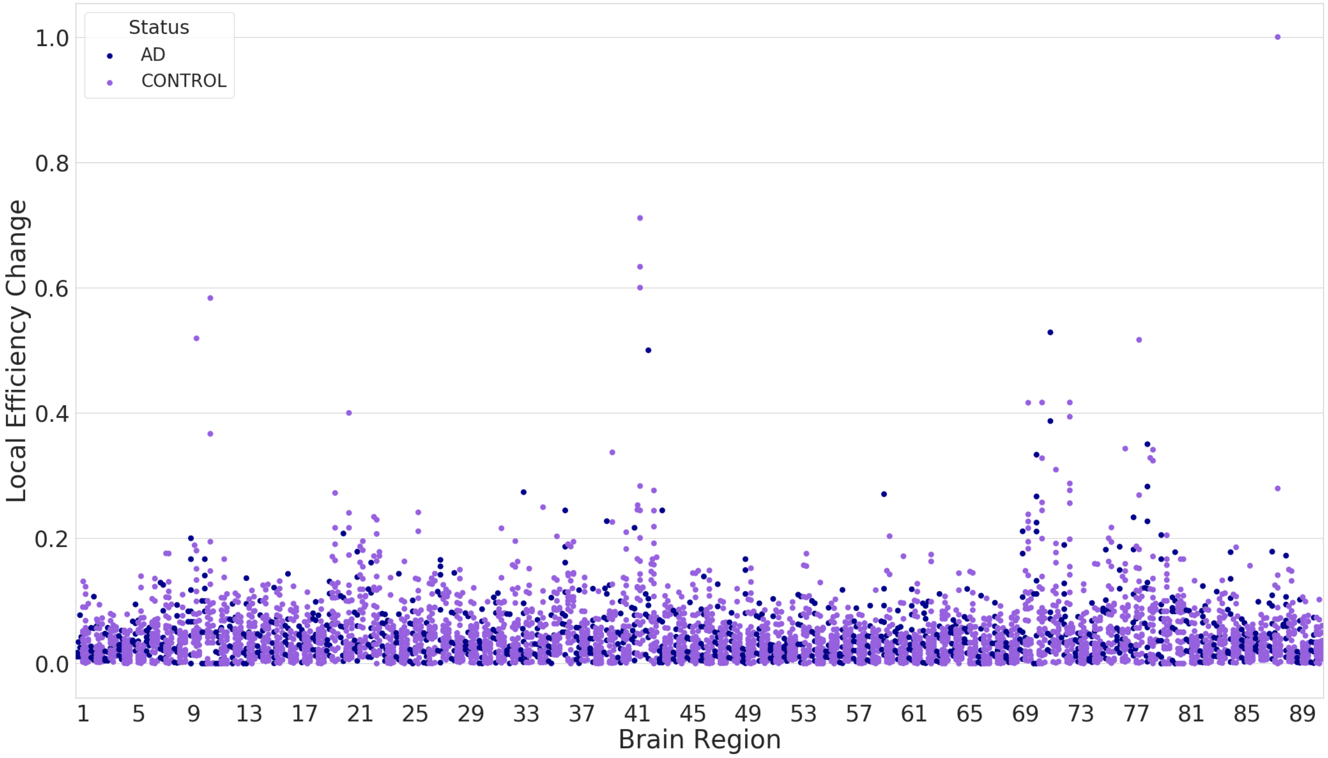


**Figure S1.** The figure shows the distribution of the absolute differences between the baseline and follow-up measures of local efficiency of the AD (blue) vs controls (purple), along the 90 AAL brain regions.

**Table S1.** Full names of brain AAL atlas regions.

| Region acronym | Region name | Abbr. | Region id |
|---|---|---|---|
| **Precentral_L** | Precental gyrus | PreCG.L | 1 |
| **Precentral_R** | Precental gyrus | PreCG.R | 2 |
| **Frontal_Sup_L** | Superior frontal gyrus;dorsolateral | SFGdor.L | 3 |
| **Frontal_Sup_R** | Superior frontal gyrus;dorsolateral | SFGdor.R | 4 |
| **Frontal_Sup_Orb_L** | Superior frontal gyrus; orbital part | ORBsup.L | 5 |
| **Frontal_Sup_Orb_R** | Superior frontal gyrus; orbital part | ORBsup.R | 6 |
| **Frontal_Mid_L** | Middle frontal gyrus | MFG.L | 7 |
| **Frontal_Mid_R** | Middle frontal gyrus | MFG.R | 8 |
| **Frontal_Mid_Orb_L** | Middle frontal gyrus; orbital part | ORBmid.L | 9 |
| **Frontal_Mid_Orb_R** | Middle frontal gyrus; orbital part | ORBmid.R | 10 |
| **Frontal_Inf_Oper_L** | Inferior frontal gyrus;opercular part | IFGoperc.L | 11 |
| **Frontal_Inf_Oper_R** | Inferior frontal gyrus;opercular part | IFGoperc.R | 12 |
| **Frontal_Inf_Tri_L** | Inferior frontal gyrus;triangular part | IFGtriang.L | 13 |
| **Frontal_Inf_Tri_R** | Inferior frontal gyrus;triangular part | IFGtriang.R | 14 |
| **Frontal_Inf_Orb_L** | Inferior frontal gyrus; orbitalpart | ORBinf.L | 15 |
| **Frontal_Inf_Orb_R** | Inferior frontal gyrus; orbitalpart | ORBinf.R | 16 |
| **Rolandic_Oper_L** | Rolandic operculum | ROL.L | 17 |
| **Rolandic_Oper_R** | Rolandic operculum | ROL.R | 18 |
| **Supp_Motor_Area_L** | Supplementary motor area | SMA.L | 19 |
| **Supp_Motor_Area_R** | Supplementary motor area | SMA.R | 20 |
| **Olfactory_L** | Olfactory cortex | OLF.L | 21 |
| **Olfactory_R** | Olfactory cortex | OLF.R | 22 |
| **Frontal_Sup_Medial_L** | Superior frontal gyrus; medial | SFGmed.L | 23 |
| **Frontal_Sup_Medial_R** | Superior frontal gyrus; medial | SFGmed.R | 24 |
| **Frontal_Mid_Orb_L** | Superior frontal gyrus; medial orbital | ORBsupmed.L | 25 |
| **Frontal_Mid_Orb_R** | Superior frontal gyrus; medial orbital | ORBsupmed.R | 26 |
| **Rectus_L** | Gyrus rectus | REC.L | 27 |
| **Rectus_R** | Gyrus rectus | REC.R | 28 |
| **Insula_L** | Insula | INS.L | 29 |
| **Insula_R** | Insula | INS.R | 30 |
| **Cingulum_Ant_L** | Anterior cingulate and paracingulate gyri | ACG.L | 31 |
| **Cingulum_Ant_R** | Anterior cingulate and paracingulate gyri | ACG.R | 32 |
| **Cingulum_Mid_L** | Median cingulate and paracingulate gyri | DCG.L | 33 |
| **Cingulum_Mid_R** | Median cingulate and paracingulate gyri | DCG.R | 34 |
| **Cingulum_Post_L** | Posterior cingulate gyrus | PCG.L | 35 |
| **Cingulum_Post_R** | Posterior cingulate gyrus | PCG.R | 36 |
| **Hippocampus_L** | Hippocampus | HIP.L | 37 |
| **Hippocampus_R** | Hippocampus | HIP.R | 38 |
| **ParaHippocampal_L** | Parahippocampal gyrus | PHG.L | 39 |
| **ParaHippocampal_R** | Parahippocampal gyrus | PHG.R | 40 |
| **Amygdala_L** | Amygdala | AMYG.L | 41 |
| **Amygdala_R** | Amygdala | AMYG.R | 42 |
| **Calcarine_L** | Calcarine fissure and surrounding cortex | CAL.L | 43 |
| **Calcarine_R** | Calcarine fissure and surrounding cortex | CAL.R | 44 |
| **Cuneus_L** | Cuneus | CUN.L | 45 |
| **Cuneus_R** | Cuneus | CUN.R | 46 |
| **Lingual_L** | Lingual gyrus | LING.L | 47 |
| **Lingual_R** | Lingual gyrus | LING.R | 48 |
| **Occipital_Sup_L** | Superior occipital gyrus | SOG.L | 49 |
| **Occipital_Sup_R** | Superior occipital gyrus | SOG.R | 50 |

**Table S1.** Full names of brain AAL atlas regions (continued).

| Region acronym | Region name | Abbr. | Region id |
|---|---|---|---|
| **Occipital_Mid_L** | Middle occipital gyrus | MOG.L | 51 |
| **Occipital_Mid_R** | Middle occipital gyrus | MOG.R | 52 |
| **Occipital_Inf_L** | Inferior occipital gyrus | IOG.L | 53 |
| **Occipital_Inf_R** | Inferior occipital gyrus | IOG.R | 54 |
| **Fusiform_L** | Fusiform gyrus | FFG.L | 55 |
| **Fusiform_R** | Fusiform gyrus | FFG.R | 56 |
| **Postcentral_L** | Postcentral gyrus | PoCG.L | 57 |
| **Postcentral_R** | Postcentral gyrus | PoCG.R | 58 |
| **Parietal_Sup_L** | Superior parietal gyrus | SPG.L | 59 |
| **Parietal_Sup_R** | Superior parietal gyrus | SPG.R | 60 |
| **Parietal_Inf_L** | Inferior parietal; but supramarginal and angular gyri | IPL.L | 61 |
| **Parietal_Inf_R** | Inferior parietal; but supramarginal and angular gyri | IPL.R | 62 |
| **SupraMarginal_L** | Supramarginal gyrus | SMG.L | 63 |
| **SupraMarginal_R** | Supramarginal gyrus | SMG.R | 64 |
| **Angular_L** | Angular gyrus | ANG.L | 65 |
| **Angular_R** | Angular gyrus | ANG.R | 66 |
| **Precuneus_L** | Precuneus | PCUN.L | 67 |
| **Precuneus_R** | Precuneus | PCUN.R | 68 |
| **Paracentral_Lobule_L** | Paracentral lobule | PCL.L | 69 |
| **Paracentral_Lobule_R** | Paracentral lobule | PCL.R | 70 |
| **Caudate_L** | Caudate nucleus | CAU.L | 71 |
| **Caudate_R** | Caudate nucleus | CAU.R | 72 |
| **Putamen_L** | Lenticular nucleus; putamen | PUT.L | 73 |
| **Putamen_R** | Lenticular nucleus; putamen | PUT.R | 74 |
| **Pallidum_L** | Lenticular nucleus; pallidum | PAL.L | 75 |
| **Pallidum_R** | Lenticular nucleus; pallidum | PAL.R | 76 |
| **Thalamus_L** | Thalamus | THA.L | 77 |
| **Thalamus_R** | Thalamus | THA.R | 78 |
| **Heschl_L** | Heschl gyrus | HES.L | 79 |
| **Heschl_R** | Heschl gyrus | HES.R | 80 |
| **Temporal_Sup_L** | Superior temporal gyrus | STG.L | 81 |
| **Temporal_Sup_R** | Superior temporal gyrus | STG.R | 82 |
| **Temporal_Pole_Sup_L** | Temporal pole: superior temporal gyrus | TPOsup.L | 83 |
| **Temporal_Pole_Sup_R** | Temporal pole: superior temporal gyrus | TPOsup.R | 84 |
| **Temporal_Mid_L** | Middle temporal gyrus | MTG.L | 85 |
| **Temporal_Mid_R** | Middle temporal gyrus | MTG.R | 86 |
| **Temporal_Pole_Mid_L** | Temporal pole: middle temporal gyrus | TPOmid.L | 87 |
| **Temporal_Pole_Mid_R** | Temporal pole: middle temporal gyrus | TPOmid.R | 88 |
| **Temporal_Inf_L** | Inferior temporal gyrus | ITG.L | 89 |
| **Temporal_Inf_R** | Inferior temporal gyrus | ITG.R | 90 |

**Table S2.** Quantile regression top results of regressing CDR scores on the local connectivity metrics.

| **CDR** | **Results are sorted according to p-value. Threshold**= $\frac{0.05}{6\times 90} = 9.26e-05$ | | | | |
|---|---|---|---|---|---|
| | Metric | Region | Region id | $\beta$ | P-value |
| **CDJUDGE** | betweencentrality | Frontal_Inf_Oper_L | 11 | -1.06e-08 | 1.3246e-17 |
| **CDCOMMUN** | betweencentrality | Frontal_Inf_Tri_L | 13 | 1.162e-07 | 1.0377e-16 |
| **CDCOMMUN** | betweencentrality | Pallidum_R | 76 | 6.79e-08 | 1.5932e-16 |
| **CDCARE** | betweencentrality | Pallidum_R | 76 | 1.21e-08 | 2.5409e-15 |
| **CDCARE** | betweencentrality | Frontal_Inf_Tri_L | 13 | -2.35e-08 | 4.3817e-15 |
| **CDCARE** | betweencentrality | Rolandic_Oper_R | 18 | -5e-09 | 5.5180e-14 |
| **CDCARE** | betweencentrality | Frontal_Mid_Orb_L | 9 | 8.35e-08 | 6.8455e-14 |
| **CDCARE** | betweencentrality | Frontal_Inf_Tri_R | 14 | 1.538e-07 | 4.6868e-13 |
| **CDMEMORY** | betweencentrality | Pallidum_R | 76 | 9.8e-09 | 1.8588e-12 |
| **CDMEMORY** | betweencentrality | Heschl_R | 80 | -1.431e-07 | 7.9135e-12 |
| **CDHOME** | betweencentrality | Pallidum_R | 76 | 2.34e-08 | 1.0339e-10 |
| **CDORIENT** | betweencentrality | Rolandic_Oper_R | 18 | 3.566e-07 | 2.8690e-10 |
| **CDCOMMUN** | betweencentrality | Frontal_Sup_Medial_L | 23 | 1.2e-08 | 2.2249e-09 |
| **CDMEMORY** | betweencentrality | Frontal_Mid_Orb_L | 9 | 1.53e-08 | 2.7792e-09 |
| **CDHOME** | betweencentrality | Frontal_Inf_Tri_L | 13 | 7.55e-08 | 2.8144e-09 |
| **CDCARE** | local_eff | Parietal_Sup_L | 59 | -2.8362e-06 | 4.5150e-09 |
| **CDCARE** | betweencentrality | Caudate_R | 72 | -2.7e-09 | 5.9383e-08 |
| **CDCARE** | betweencentrality | Frontal_Sup_Medial_L | 23 | 1.81e-08 | 8.4091e-08 |
| **CDCOMMUN** | betweencentrality | Precentral_L | 1 | 5.6e-09 | 8.8635e-08 |
| **CDCARE** | betweencentrality | Frontal_Sup_Orb_L | 5 | -6.8e-09 | 9.1463e-08 |
| **CDCARE** | betweencentrality | Occipital_Inf_L | 53 | 3.2e-09 | 1.0397e-07 |
| **CDJUDGE** | betweencentrality | Insula_R | 30 | -1.9e-09 | 1.0927e-07 |
| **CDORIENT** | betweencentrality | Angular_R | 66 | -2.58e-08 | 1.3192e-07 |
| **CDHOME** | betweencentrality | Frontal_Sup_Medial_L | 23 | 7.28e-08 | 1.4609e-07 |
| **CDMEMORY** | betweencentrality | Angular_L | 65 | 9.69e-08 | 1.5599e-07 |
| **CDCOMMUN** | betweencentrality | Occipital_Inf_L | 53 | 8e-09 | 1.6902e-07 |
| **CDCOMMUN** | betweencentrality | Occipital_Sup_R | 50 | 8.8e-09 | 1.8204e-07 |
| **CDCARE** | betweencentrality | Cingulum_Post_R | 36 | 3.55e-08 | 2.6603e-07 |
| **CDCARE** | betweencentrality | Angular_L | 65 | 5.9e-09 | 3.2401e-07 |
| **CDCARE** | betweencentrality | Frontal_Inf_Oper_R | 12 | -4.9e-09 | 3.9313e-07 |
| **CDMEMORY** | betweencentrality | Frontal_Inf_Oper_R | 12 | 1.72e-08 | 4.2464e-07 |
| **CDJUDGE** | betweencentrality | Precentral_R | 2 | 2.4e-09 | 4.2576e-07 |
| **CDJUDGE** | betweencentrality | Paracentral_Lobule_R | 70 | 5.1e-09 | 4.5211e-07 |
| **CDCARE** | local_eff | Caudate_L | 71 | 7.203e-07 | 6.0570e-07 |
| **CDMEMORY** | betweencentrality | Occipital_Mid_R | 52 | -1.45e-08 | 7.3061e-07 |
| **CDJUDGE** | betweencentrality | SupraMarginal_R | 64 | -1.35e-08 | 7.3933e-07 |
| **CDJUDGE** | betweencentrality | Calcarine_L | 43 | 1.2e-09 | 8.0399e-07 |
| **CDCARE** | betweencentrality | Precentral_L | 1 | 3.9e-09 | 1.0015e-06 |
| **CDCARE** | local_eff | Thalamus_R | 78 | -1.6719e-06 | 1.1912e-06 |
| **CDCARE** | betweencentrality | ParaHippocampal_L | 39 | 8.1e-09 | 1.2821e-06 |
| **CDMEMORY** | betweencentrality | Precentral_L | 1 | 4.9e-09 | 1.4549e-06 |
| **CDJUDGE** | betweencentrality | Cingulum_Mid_L | 33 | 5e-10 | 1.8206e-06 |
| **CDORIENT** | betweencentrality | Amygdala_R | 42 | 2.7e-08 | 2.0376e-06 |
| **CDCARE** | betweencentrality | Cuneus_R | 46 | 2.6e-09 | 2.1374e-06 |
| **CDJUDGE** | local_eff | Occipital_Mid_L | 51 | -1.3593e-06 | 2.5568e-06 |
| **CDCARE** | betweencentrality | Occipital_Sup_R | 50 | 2.4e-09 | 2.7811e-06 |
| **CDCARE** | betweencentrality | SupraMarginal_R | 64 | 2.35e-08 | 2.8475e-06 |
| **CDCARE** | cluster_coef | Parietal_Sup_L | 59 | -1.5812e-06 | 3.0162e-06 |
| **CDCARE** | betweencentrality | Precentral_R | 2 | 6.4e-09 | 3.0711e-06 |
| **CDJUDGE** | betweencentrality | Cuneus_R | 46 | 8e-10 | 3.1560e-06 |

**Table S2.** Quantile regression top results of regressing CDR scores on the local connectivity metrics (continued).

| CDR | Results are sorted according to p-value. Threshold= $\frac{0.05}{6\times 90} = 9.26e-05$ | | | | |
|---|---|---|---|---|---|
| | Metric | Region | Region id | $\beta$ | P-value |
| **CDMEMORY** | betweencentrality | Precentral_R | 2 | 1.21e-08 | 4.0628e-06 |
| **CDCARE** | betweencentrality | Occipital_Mid_L | 51 | -2e-09 | 4.7259e-06 |
| **CDCARE** | betweencentrality | Temporal_Inf_R | 90 | -8e-10 | 5.1879e-06 |
| **CDCOMMUN** | betweencentrality | Temporal_Pole_Sup_R | 84 | 1.7e-09 | 5.2490e-06 |
| **CDCARE** | local_eff | Paracentral_Lobule_R | 70 | 4.028e-07 | 5.3093e-06 |
| **CDCARE** | betweencentrality | Olfactory_R | 22 | 2.5e-09 | 6.1963e-06 |
| **CDCARE** | betweencentrality | Pallidum_L | 75 | 2.8e-09 | 6.6154e-06 |
| **CDJUDGE** | betweencentrality | Postcentral_L | 57 | -4e-10 | 6.6330e-06 |
| **CDCARE** | betweencentrality | Frontal_Med_Orb_L | 25 | -5.5e-09 | 6.7257e-06 |
| **CDCARE** | betweencentrality | Parietal_Inf_L | 61 | -7.6e-09 | 6.8700e-06 |
| **CDCARE** | local_eff | Calcarine_L | 43 | -1.3701e-06 | 6.9599e-06 |
| **CDCARE** | betweencentrality | Cingulum_Mid_L | 33 | 9e-10 | 7.0795e-06 |
| **CDHOME** | betweencentrality | Precentral_L | 1 | 6.5e-09 | 7.2322e-06 |
| **CDJUDGE** | cluster_coef | Occipital_Mid_L | 51 | -7.789e-07 | 7.5340e-06 |
| **CDCARE** | betweencentrality | Olfactory_L | 21 | 1.2e-09 | 1.0177e-05 |
| **CDCARE** | betweencentrality | Paracentral_Lobule_L | 69 | 2.9e-09 | 1.0179e-05 |
| **CDORIENT** | betweencentrality | Frontal_Sup_Medial_L | 23 | 4.18e-08 | 1.1987e-05 |
| **CDCARE** | cluster_coef | Thalamus_R | 78 | -9.55e-07 | 1.2007e-05 |
| **CDMEMORY** | betweencentrality | Frontal_Sup_Orb_L | 5 | -2.3e-09 | 1.3418e-05 |
| **CDCARE** | betweencentrality | Frontal_Sup_Medial_R | 24 | -2.2e-09 | 1.3753e-05 |
| **CDMEMORY** | betweencentrality | Thalamus_L | 77 | 4.6e-09 | 1.4079e-05 |
| **CDCARE** | betweencentrality | Putamen_R | 74 | -6.6e-09 | 1.4768e-05 |
| **CDCOMMUN** | betweencentrality | Putamen_R | 74 | -1.1e-09 | 1.7248e-05 |
| **CDCARE** | local_eff | Amygdala_R | 42 | -1.2503e-06 | 1.7740e-05 |
| **CDCARE** | betweencentrality | Thalamus_L | 77 | -7e-10 | 1.7820e-05 |
| **CDJUDGE** | betweencentrality | Frontal_Inf_Orb_R | 16 | 6e-10 | 1.7857e-05 |
| **CDJUDGE** | cluster_coef | Temporal_Mid_L | 85 | 6.814e-07 | 1.7954e-05 |
| **CDJUDGE** | betweencentrality | Pallidum_L | 75 | 6e-10 | 1.9036e-05 |
| **CDCARE** | local_eff | Lingual_L | 47 | -1.3648e-06 | 1.9534e-05 |
| **CDCARE** | betweencentrality | Putamen_L | 73 | -9e-10 | 2.0543e-05 |
| **CDCARE** | local_eff | Frontal_Mid_Orb_L | 9 | -9.801e-07 | 2.0693e-05 |
| **CDJUDGE** | local_eff | Temporal_Mid_L | 85 | 1.1769e-06 | 2.4200e-05 |
| **CDJUDGE** | cluster_coef | Frontal_Sup_Orb_L | 5 | 5.54e-07 | 2.5902e-05 |
| **CDCARE** | betweencentrality | Temporal_Pole_Sup_L | 83 | -2.4e-09 | 2.7526e-05 |
| **CDJUDGE** | local_eff | Calcarine_L | 43 | -5.122e-07 | 2.7953e-05 |
| **CDJUDGE** | cluster_coef | Cuneus_R | 46 | -6.206e-07 | 2.8360e-05 |
| **CDCARE** | betweencentrality | Frontal_Med_Orb_R | 26 | 1.1e-09 | 2.9931e-05 |
| **CDCARE** | betweencentrality | Rectus_L | 27 | 1.7e-09 | 3.1086e-05 |
| **CDCARE** | betweencentrality | Temporal_Pole_Sup_R | 84 | 4e-10 | 3.2473e-05 |
| **CDCARE** | local_eff | Precentral_R | 2 | -1.7287e-06 | 3.3132e-05 |
| **CDJUDGE** | local_eff | Frontal_Sup_Orb_L | 5 | 7.069e-07 | 3.3295e-05 |
| **CDCARE** | cluster_coef | Precuneus_L | 67 | -1.3847e-06 | 3.3801e-05 |
| **CDJUDGE** | cluster_coef | Olfactory_L | 21 | 3.541e-07 | 3.5295e-05 |
| **CDCARE** | cluster_coef | Occipital_Sup_L | 49 | -1.2676e-06 | 3.5428e-05 |
| **CDJUDGE** | cluster_coef | Calcarine_L | 43 | -4.54e-07 | 3.7033e-05 |
| **CDJUDGE** | local_eff | Cuneus_R | 46 | -1.0686e-06 | 3.7315e-05 |
| **CDCARE** | local_eff | Occipital_Sup_L | 49 | -2.3923e-06 | 3.9588e-05 |
| **CDCOMMUN** | betweencentrality | Frontal_Sup_Orb_R | 6 | 1e-09 | 4.0230e-05 |
| **CDCARE** | betweencentrality | Rolandic_Oper_L | 17 | 6.2e-08 | 4.0397e-05 |

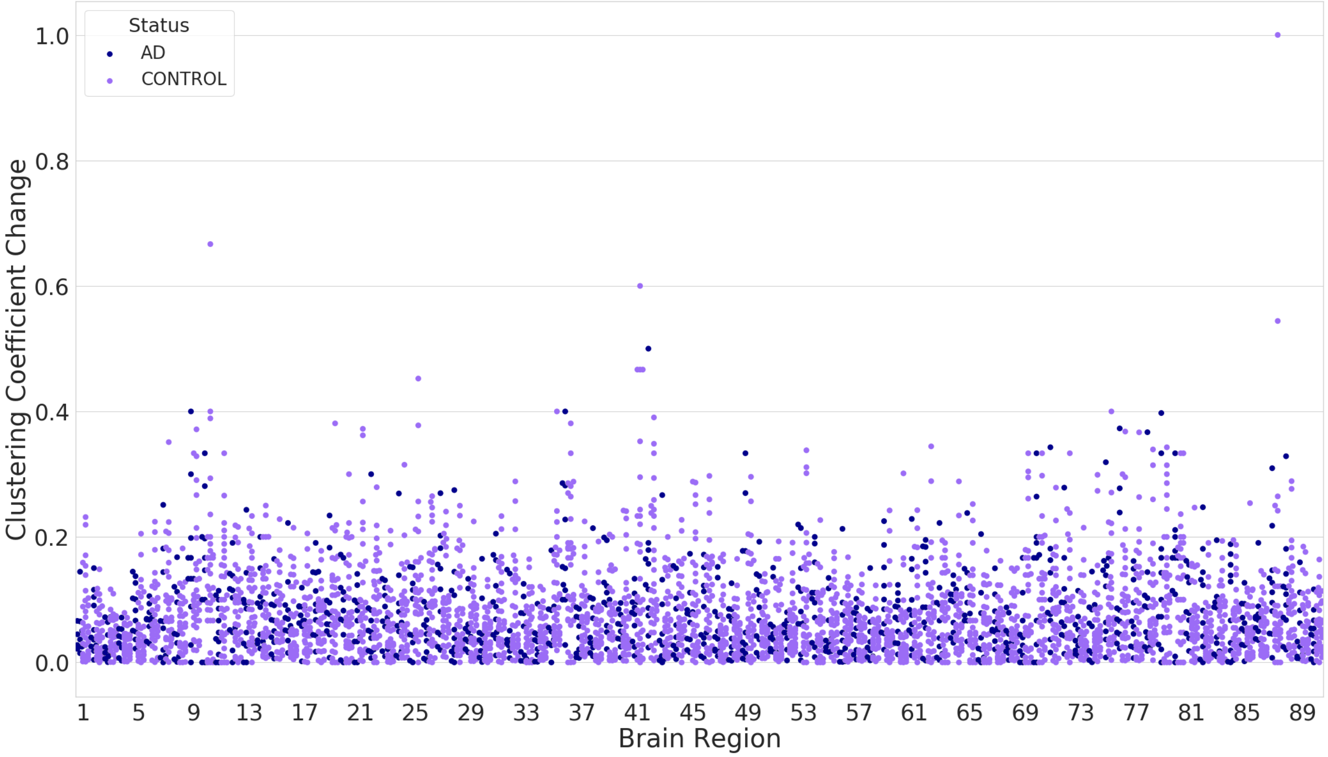


**Figure S2.** The figure shows the distribution of the absolute differences between the baseline and follow-up measures of clustering coefficient of the AD (blue) vs controls (purple), along the 90 AAL brain regions.

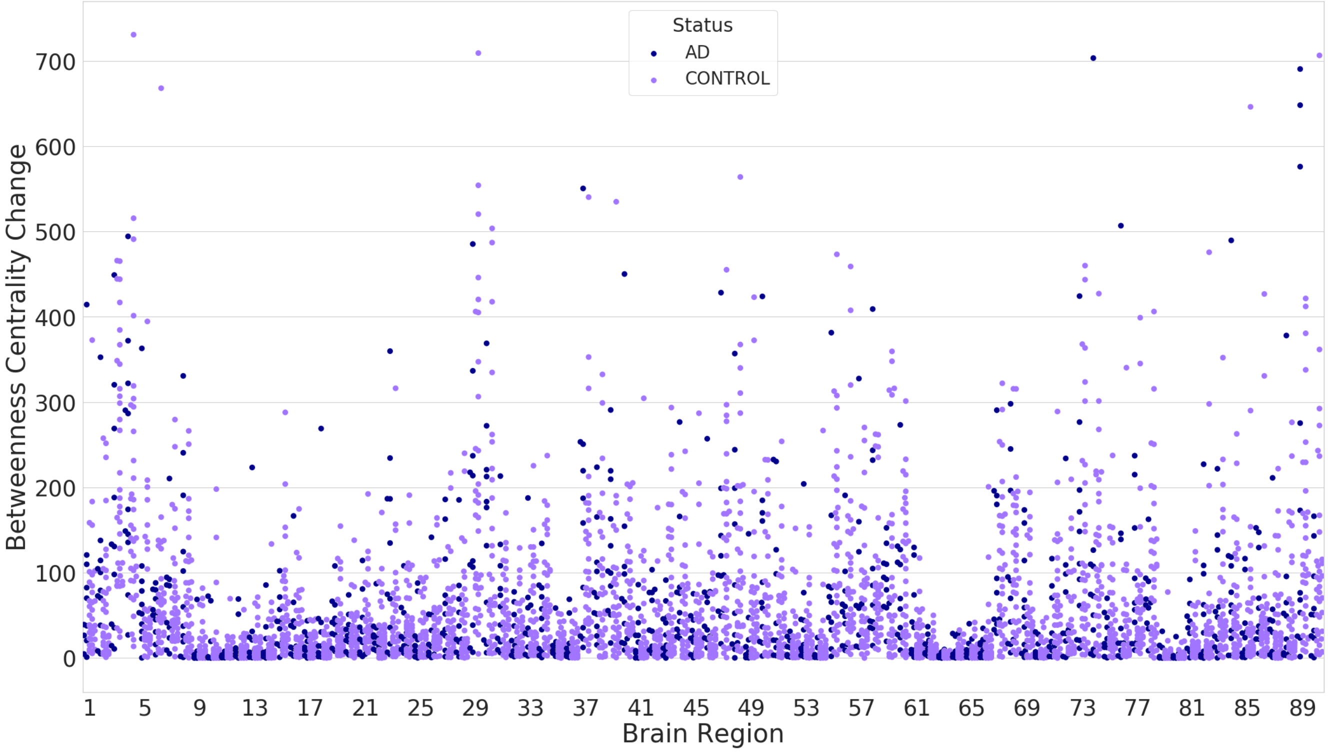


**Figure S3.** The figure shows the distribution of the absolute differences between the baseline and follow-up measures of betweenness centrality of the AD (blue) vs controls (purple), along the 90 AAL brain regions.

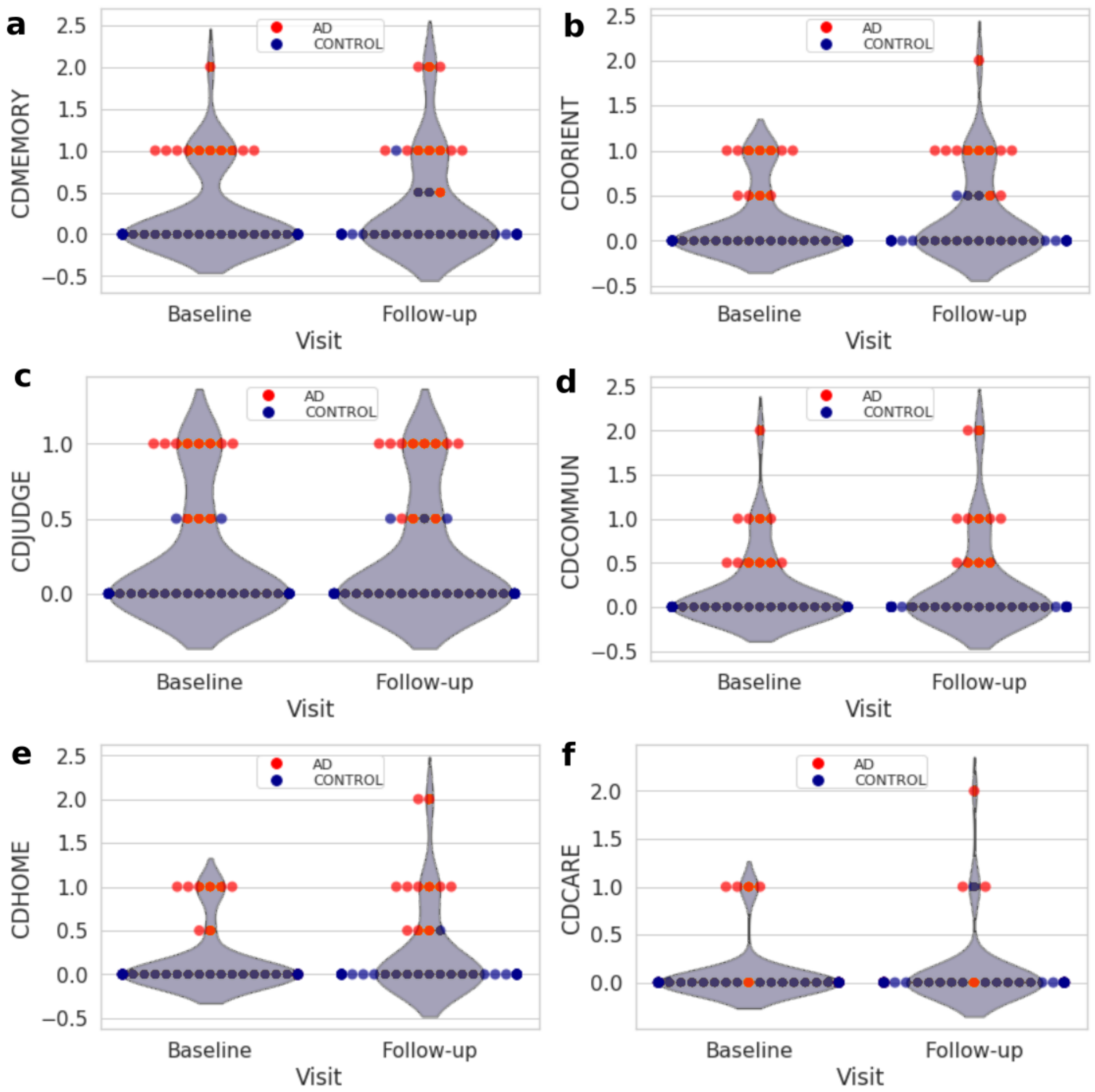


**Figure S4.** Violin plots to illustrate the CDR scores (either 0: None, 0.5: very mild, 1: mild, 2: moderate or 3: severe) in the baseline (left violin plot) and follow-up (right violin plot) visits, for AD (red dots) and controls (blue dots). The more dots moved to the higher scores from Baseline to Follow-up, the more patients worsened their disabilities. The memory (CDMEMORY; a) and orientation (CDORIENT; b) scores are represented by the top sub-figures, judgment and problem solving (CDJUDGE; c) and community affairs (CDCOMMUN; d) are the middle sub-figures, while home and hobbies (CDHOME; e) and personal care (CDCARE; f) are at the bottom. It is visible that generally some AD subjects worsen their score.

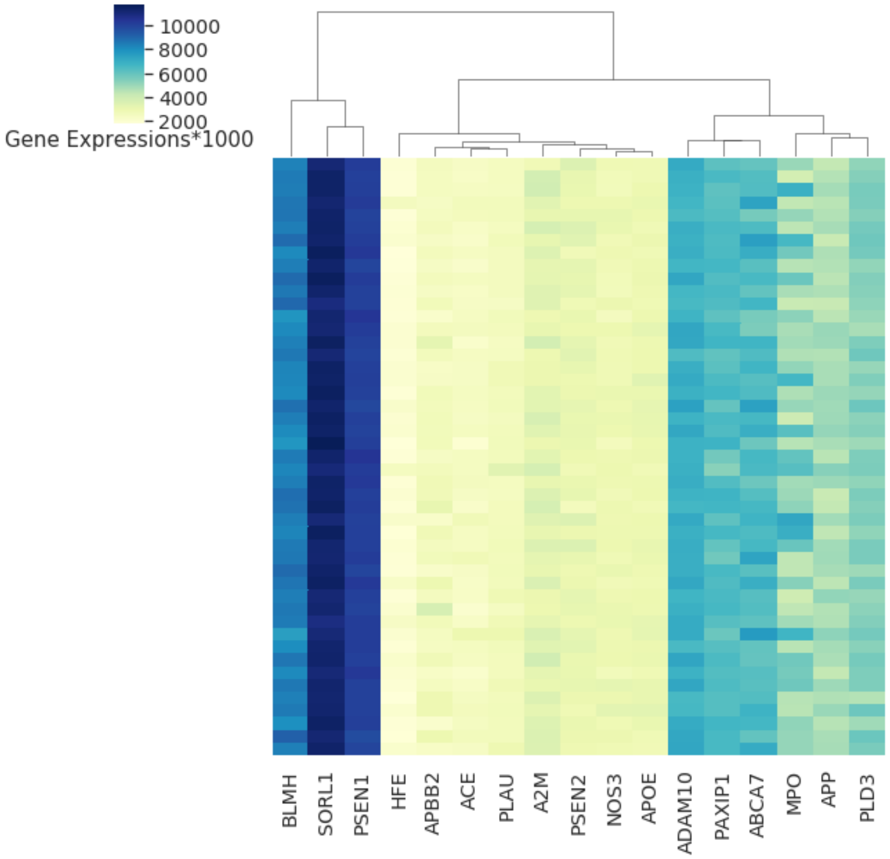


**Figure S5.** A heatmap of the estimated 17 gene expression profiles (values multiplied by 1000, each line represents a participant) out of the 65 probe sets as explained in the Materials and Methods section. The dark blue represents a high expression values, while the yellow represents low expression. The SORL1 has the highest expression among the genes and HFE expression was the lowest among other genes.

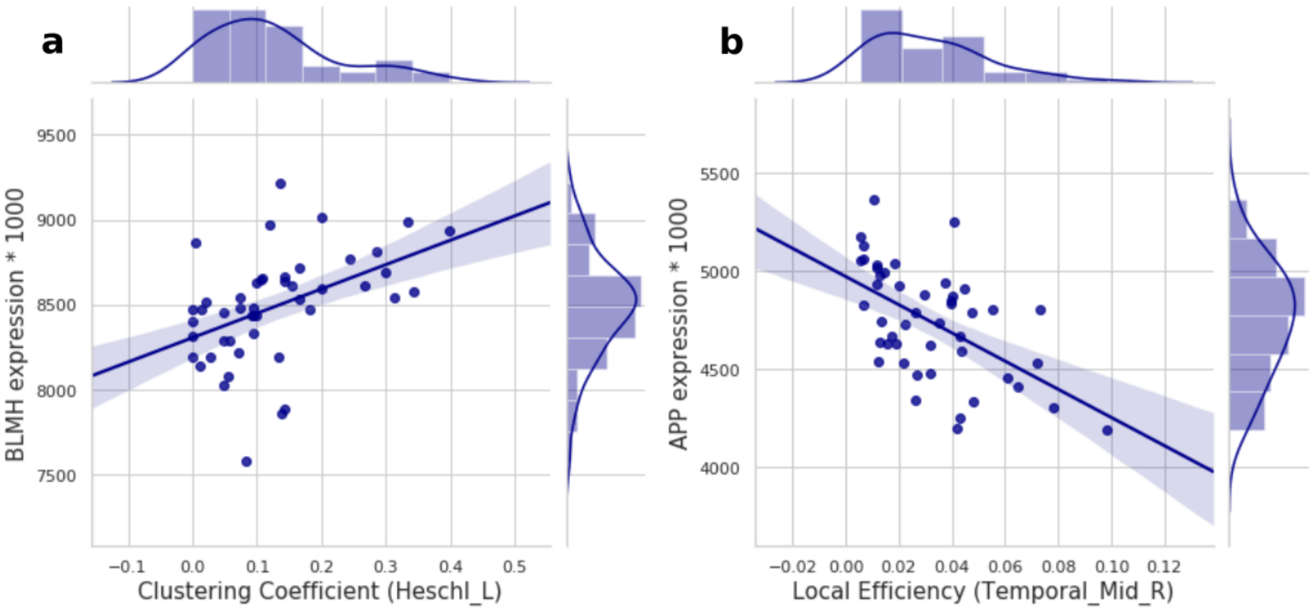


**Figure S6.** A scatter plot of all the significant association results. The plots shows the associations between; (a) BLMH expression and clustering coefficient in AAL region 79 (Heschl_L), (b) APP expression and local efficiency in brain region 86 (Temporal_Mid_R).

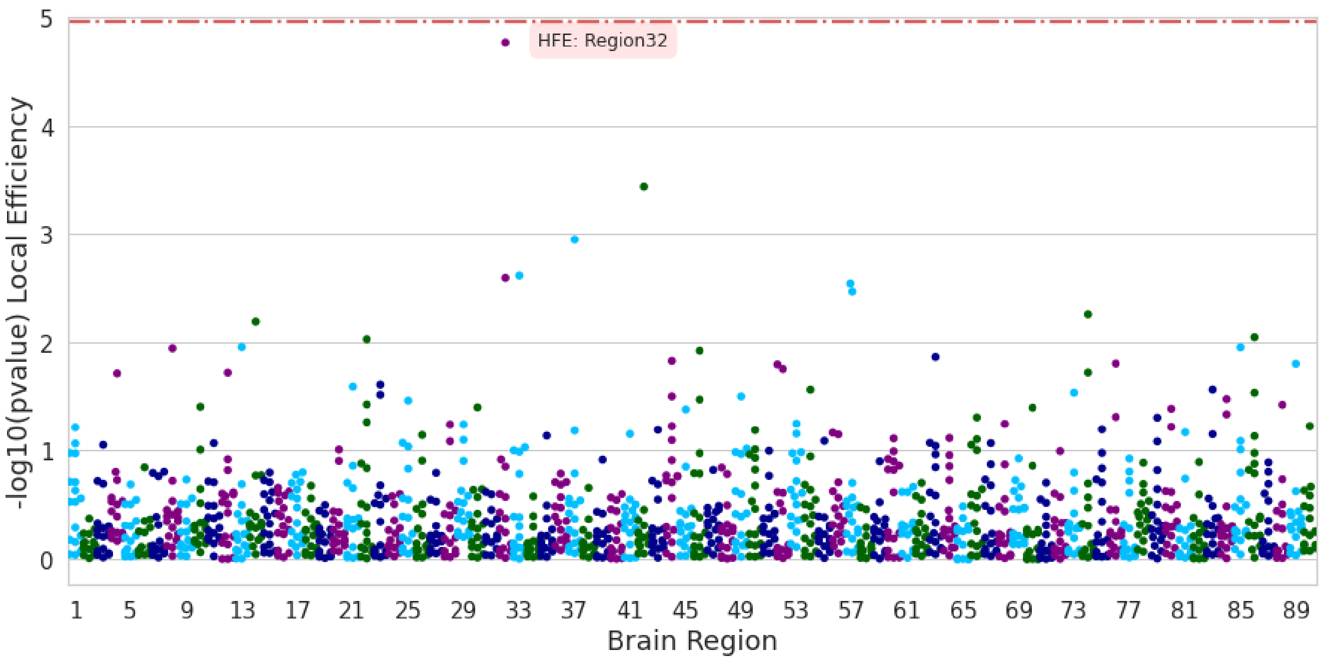


**Figure S7.** The figure shows the quantile regression model coefficient $-\log 10$p-values. The model regresses the change in the local coefficient (dependant variable) on a single gene at a time (independent variable), at each of the 90 brain regions as in the AAL atlas (x axis).

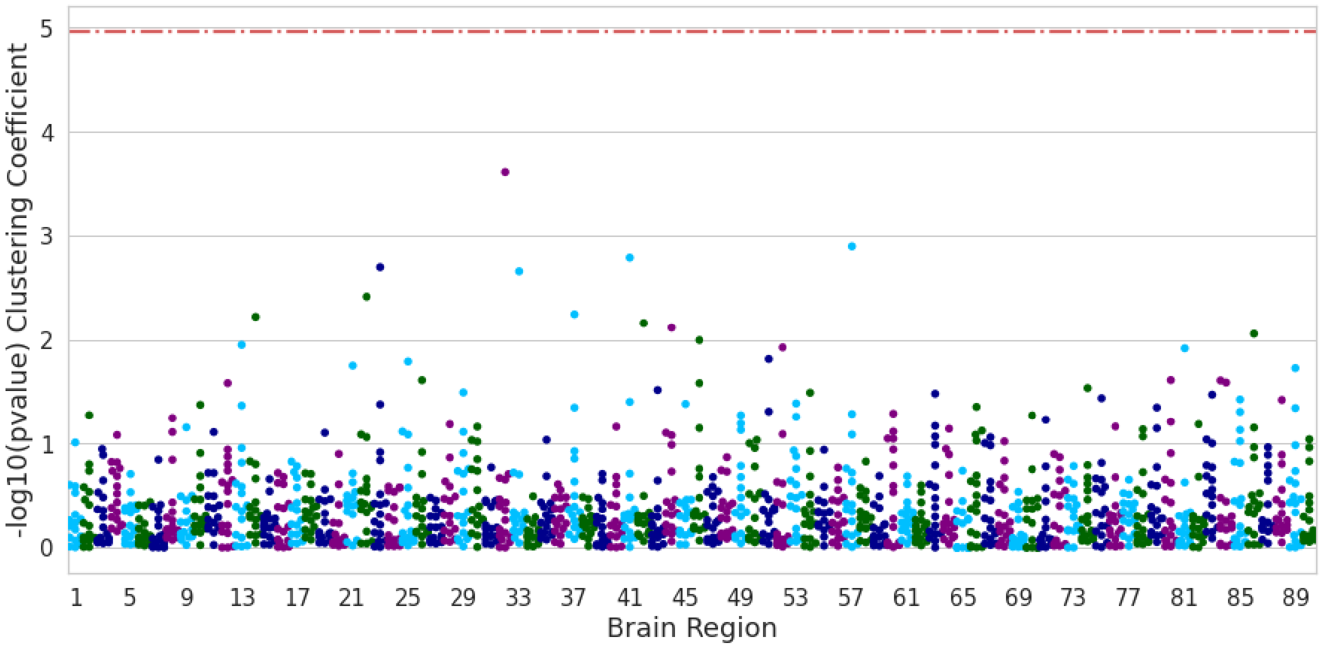


**Figure S8.** The figure shows the quantile regression model coefficient $-\log 10$p-values. The model regresses the change in the betweenness centrality (dependant variable) on a single gene at a time (independent variable), at each of the 90 brain regions as in the AAL atlas (x axis).

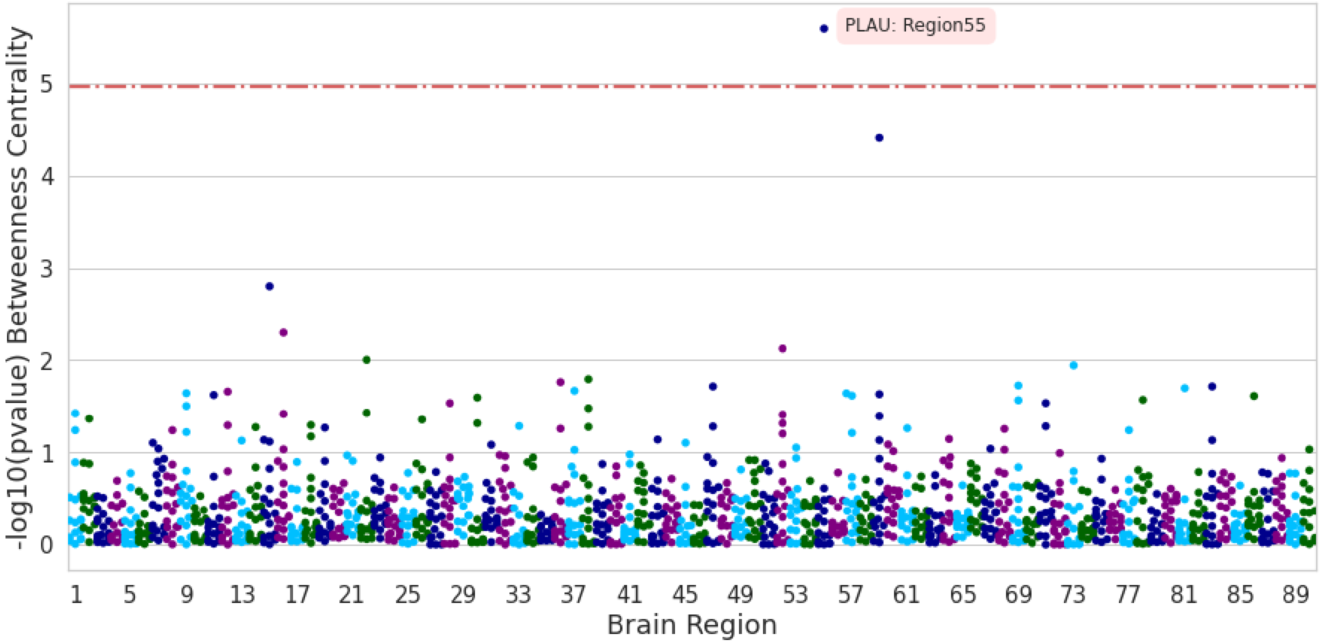


**Figure S9.** The figure shows the quantile regression model coefficient $-\log 10$p-values. The model regresses the change in the local coefficient (dependant variable) on a single gene at a time (independent variable), at each of the 90 brain regions as in the AAL atlas (x axis).

**Table S2.** Quantile regression top results of regressing CDR scores on the local connectivity metrics (continued).

| **CDR** | **Results are sorted according to p-value. Threshold=** $\frac{0.05}{6\times90} = 9.26e-05$ | | | | |
|---|---|---|---|---|---|
| | Metric | Region | Region id | $\beta$ | P-value |
| **CDJUDGE** | betweencentrality | Insula_L | 29 | 2e-10 | 4.3809e-05 |
| **CDR_diff** | betweencentrality | Frontal_Sup_Medial_L | 23 | 0.0037021312 | 4.5916e-05 |
| **CDCOMMUN** | cluster_coef | Precuneus_L | 67 | -1.9329e-06 | 4.9971e-05 |
| **CDCARE** | local_eff | Occipital_Mid_L | 51 | -1.4926e-06 | 5.3963e-05 |
| **CDCARE** | betweencentrality | Temporal_Inf_L | 89 | 2e-10 | 5.5267e-05 |
| **CDCARE** | betweencentrality | Amygdala_L | 41 | -1.1e-09 | 5.6693e-05 |
| **CDCARE** | betweencentrality | Frontal_Inf_Orb_L | 15 | -8e-10 | 5.6717e-05 |
| **CDJUDGE** | betweencentrality | Putamen_R | 74 | 3e-10 | 5.8176e-05 |
| **CDCARE** | local_eff | Parietal_Inf_L | 61 | 1.8772e-06 | 5.9063e-05 |
| **CDORIENT** | local_eff | Caudate_L | 71 | 2.6838e-06 | 5.9186e-05 |
| **CDCARE** | local_eff | ParaHippocampal_L | 39 | -6.989e-07 | 6.0628e-05 |
| **CDCARE** | cluster_coef | Paracentral_Lobule_L | 69 | -3.682e-07 | 6.0769e-05 |
| **CDHOME** | betweencentrality | Cingulum_Mid_L | 33 | 1.12e-08 | 6.2336e-05 |
| **CDCARE** | local_eff | Temporal_Pole_Sup_R | 84 | -2.5008e-06 | 6.4671e-05 |
| **CDCARE** | betweencentrality | Calcarine_R | 44 | -2.3e-09 | 7.4457e-05 |
| **CDCARE** | betweencentrality | ParaHippocampal_R | 40 | -1.2e-09 | 7.5724e-05 |
| **CDMEMORY** | local_eff | Temporal_Mid_L | 85 | 4.7978e-06 | 7.5984e-05 |
| **CDORIENT** | betweencentrality | Thalamus_L | 77 | 3.7e-09 | 8.3093e-05 |
| **CDCARE** | cluster_coef | Parietal_Inf_L | 61 | 9.696e-07 | 8.3094e-05 |
| **CDCARE** | betweencentrality | Precuneus_L | 67 | 1.3e-09 | 8.9269e-05 |
| **CDJUDGE** | local_eff | Precentral_R | 2 | -1.0795e-06 | 8.9400e-05 |
| **CDCOMMUN** | betweencentrality | Cingulum_Mid_L | 33 | 1.8e-09 | 8.9822e-05 |

**Table S3.** Ridge regression results of the change CDR scores on the global connectivity changes and Alzheimer's Disease gene expressions.

| CDR | Metric | Alpha | score | Metric | *APBB2* | *MPO* | *APP* | *ACE* | *PLAU* | *PAXIP1* | *HFE* | *SORL1* | *A2M* | *NOS3* | *BLMH* | *ADAM10* | *PLD3* | *ApoE* | *PSEN1* | *PSEN2* | *ABCA7* |
|---|---|---|---|---|---|---|---|---|---|---|---|---|---|---|---|---|---|---|---|---|---|
| **CDMEMORY** | transitivity | 0.1 | -0.0747 | -0.6704 | 0.122 | 0.0823 | 0.0248 | -0.0392 | -0.0556 | 0.0497 | -0.0926 | 0.0138 | -0.3465 | 0.3276 | -0.1321 | -0.1347 | -0.0774 | -0.2426 | 0.161 | -0.1369 | 0.1903 |
| **CDMEMORY** | global_eff | 0.1 | -0.0753 | -0.0304 | 0.1114 | 0.0802 | 0.0332 | -0.0311 | -0.0575 | 0.06 | -0.0765 | 0.0086 | -0.3486 | 0.3226 | -0.1364 | -0.1318 | -0.0628 | -0.267 | 0.1585 | -0.1392 | 0.1919 |
| **CDMEMORY** | louvain | 0.1 | -0.0754 | -0.259 | 0.1052 | 0.0823 | 0.0353 | -0.0259 | -0.0676 | 0.0647 | -0.0634 | 0.0098 | -0.3483 | 0.3178 | -0.1388 | -0.1296 | -0.0616 | -0.2694 | 0.1563 | -0.1373 | 0.1884 |
| **CDMEMORY** | char_path_len | 0.1 | -0.0752 | -0.1203 | 0.1004 | 0.0804 | 0.0351 | -0.0335 | -0.0617 | 0.0619 | -0.0769 | 0.0074 | -0.3507 | 0.3187 | -0.1365 | -0.1287 | -0.0475 | -0.2827 | 0.1621 | -0.1398 | 0.1869 |
| **CDORIENT** | transitivity | 0.1 | -0.0677 | -0.3061 | 0.1731 | 0.0679 | 0.2135 | 0.0287 | 0.0046 | -0.0399 | -0.472 | 0.0156 | -0.192 | 0.0553 | -0.0138 | -0.0342 | -0.0135 | -0.1967 | 0.1309 | -0.232 | -0.0035 |
| **CDORIENT** | global_eff | 0.1 | -0.0677 | -0.0517 | 0.169 | 0.067 | 0.2172 | 0.0324 | 0.0042 | -0.0354 | -0.4648 | 0.0133 | -0.1928 | 0.0532 | -0.0157 | -0.0332 | -0.0078 | -0.2068 | 0.1297 | -0.2331 | -0.0025 |
| **CDORIENT** | louvain | 0.1 | -0.0673 | -0.403 | 0.1592 | 0.0702 | 0.2203 | 0.0406 | -0.0117 | -0.0281 | -0.4445 | 0.0151 | -0.1923 | 0.0457 | -0.0195 | -0.0298 | -0.0058 | -0.2107 | 0.1261 | -0.23 | -0.0079 |
| **CDORIENT** | char_path_len | 0.1 | -0.0677 | 0.009 | 0.1688 | 0.067 | 0.2173 | 0.0325 | 0.0039 | -0.0352 | -0.4645 | 0.0132 | -0.1929 | 0.0532 | -0.0158 | -0.033 | -0.0076 | -0.2071 | 0.1295 | -0.2331 | -0.0025 |
| **CDJUDGE** | transitivity | 149 | -0.0056 | -0.0 | -0.0002 | 0.0023 | -0.0007 | -0.0012 | 0.0 | 0.0019 | 0.0003 | 0.0002 | 0.0004 | 0.0002 | -0.0005 | -0.0002 | -0.0007 | -0.0003 | -0.0005 | -0.0001 | -0.0027 |
| **CDJUDGE** | global_eff | 149 | -0.0056 | 0.0 | -0.0002 | 0.0023 | -0.0007 | -0.0012 | 0.0 | 0.0019 | 0.0003 | 0.0002 | 0.0004 | 0.0002 | -0.0005 | -0.0002 | -0.0007 | -0.0003 | -0.0005 | -0.0001 | -0.0027 |
| **CDJUDGE** | louvain | 149 | -0.0056 | 0.0001 | -0.0002 | 0.0023 | -0.0007 | -0.0012 | 0.0 | 0.0019 | 0.0003 | 0.0002 | 0.0004 | 0.0002 | -0.0005 | -0.0002 | -0.0007 | -0.0003 | -0.0005 | -0.0001 | -0.0027 |
| **CDJUDGE** | char_path_len | 149 | -0.0056 | -0.0001 | -0.0002 | 0.0023 | -0.0007 | -0.0012 | 0.0 | 0.0019 | 0.0003 | 0.0002 | 0.0004 | 0.0002 | -0.0005 | -0.0002 | -0.0007 | -0.0003 | -0.0005 | -0.0001 | -0.0027 |
| **CDCOMMUN** | transitivity | 0.3 | -0.0325 | -0.0526 | 0.1227 | -0.0205 | -0.178 | 0.1151 | 0.1812 | -0.0161 | -0.0366 | -0.0375 | -0.1628 | 0.1476 | -0.0795 | 0.1688 | -0.0516 | 0.0563 | 0.0078 | -0.1135 | 0.0637 |
| **CDCOMMUN** | global_eff | 0.3 | -0.0325 | 0.0071 | 0.1218 | -0.0207 | -0.1775 | 0.1154 | 0.1811 | -0.0155 | -0.0356 | -0.0378 | -0.1629 | 0.1473 | -0.0798 | 0.1692 | -0.0503 | 0.0547 | 0.0076 | -0.1137 | 0.0637 |
| **CDCOMMUN** | louvain | 0.3 | -0.0325 | 0.0174 | 0.1223 | -0.0208 | -0.1776 | 0.1152 | 0.1817 | -0.0158 | -0.0363 | -0.0379 | -0.1629 | 0.1476 | -0.0797 | 0.169 | -0.0505 | 0.0548 | 0.0077 | -0.1138 | 0.064 |
| **CDCOMMUN** | char_path_len | 0.5 | -0.0324 | -0.0703 | 0.1038 | -0.0164 | -0.1436 | 0.0941 | 0.1423 | -0.0117 | -0.0184 | -0.0381 | -0.1579 | 0.1242 | -0.0666 | 0.1484 | -0.0458 | 0.0447 | 0.0139 | -0.1111 | 0.0613 |
| **CDHOME** | transitivity | 0.4 | -0.0893 | 0.0428 | 0.1908 | -0.0343 | -0.09 | 0.0756 | 0.0009 | -0.0142 | 0.0578 | -0.1228 | -0.1029 | -0.0044 | -0.1787 | 0.3326 | -0.095 | 0.1188 | 0.113 | -0.3091 | 0.1076 |
| **CDHOME** | global_eff | 0.3 | -0.0891 | 0.0872 | 0.1961 | -0.0363 | -0.0987 | 0.0821 | -0.0017 | -0.0156 | 0.0521 | -0.132 | -0.1063 | -0.0057 | -0.1846 | 0.3515 | -0.0929 | 0.1258 | 0.1227 | -0.3232 | 0.1079 |
| **CDHOME** | louvain | 0.3 | -0.0892 | -0.0191 | 0.1972 | -0.0361 | -0.0989 | 0.0826 | -0.0017 | -0.0157 | 0.0525 | -0.1318 | -0.1059 | -0.0054 | -0.1847 | 0.3511 | -0.0949 | 0.1275 | 0.1225 | -0.3231 | 0.1082 |
| **CDHOME** | char_path_len | 0.6 | -0.0891 | -0.1843 | 0.1674 | -0.0302 | -0.0767 | 0.0607 | 0.0025 | -0.0114 | 0.0616 | -0.1083 | -0.0978 | -0.0083 | -0.167 | 0.3047 | -0.0782 | 0.0932 | 0.1009 | -0.2832 | 0.0988 |
| **CDCARE** | transitivity | 149 | -0.1093 | -0.0 | -0.0008 | 0.0027 | -0.0014 | -0.0002 | 0.0007 | 0.0001 | -0.0037 | 0.0013 | -0.0066 | -0.001 | -0.0037 | -0.0018 | -0.0031 | 0.0 | 0.0013 | -0.0045 | 0.0078 |
| **CDCARE** | global_eff | 149 | -0.1093 | -0.0 | -0.0008 | 0.0027 | -0.0014 | -0.0002 | 0.0007 | 0.0001 | -0.0037 | 0.0013 | -0.0066 | -0.001 | -0.0037 | -0.0018 | -0.0031 | 0.0 | 0.0013 | -0.0045 | 0.0078 |
| **CDCARE** | louvain | 149 | -0.1093 | -0.0001 | -0.0008 | 0.0027 | -0.0014 | -0.0002 | 0.0007 | 0.0001 | -0.0037 | 0.0013 | -0.0066 | -0.001 | -0.0037 | -0.0018 | -0.0031 | 0.0 | 0.0013 | -0.0045 | 0.0078 |
| **CDCARE** | char_path_len | 149 | -0.1093 | 0.0 | -0.0008 | 0.0027 | -0.0014 | -0.0002 | 0.0007 | 0.0001 | -0.0037 | 0.0013 | -0.0066 | -0.001 | -0.0037 | -0.0018 | -0.0031 | 0.0 | 0.0013 | -0.0045 | 0.0078 |
| **CDR_diff** | transitivity | 0.1 | -1.2761 | -1.5541 | 0.7401 | 0.1734 | -0.2195 | 0.1158 | 0.197 | 0.0516 | -0.8229 | -0.0517 | -1.2177 | 0.8435 | -0.6461 | 0.3132 | -0.3519 | -0.337 | 0.6769 | -1.3473 | 0.6318 |
| **CDR_diff** | global_eff | 0.1 | -1.2782 | 0.1652 | 0.7112 | 0.1687 | -0.1988 | 0.134 | 0.1901 | 0.0768 | -0.7842 | -0.0645 | -1.2239 | 0.8302 | -0.6564 | 0.3216 | -0.3117 | -0.3999 | 0.6719 | -1.353 | 0.634 |
| **CDR_diff** | louvain | 0.1 | -1.28 | -0.4664 | 0.7041 | 0.1724 | -0.1962 | 0.1439 | 0.1741 | 0.084 | -0.762 | -0.0618 | -1.2221 | 0.823 | -0.6605 | 0.3239 | -0.3153 | -0.3984 | 0.6672 | -1.3492 | 0.6292 |
| **CDR_diff** | char_path_len | 0.1 | -1.2776 | -1.0277 | 0.6252 | 0.1706 | -0.1849 | 0.1144 | 0.1581 | 0.0904 | -0.7903 | -0.0733 | -1.2397 | 0.8002 | -0.6563 | 0.3454 | -0.1921 | -0.5225 | 0.7013 | -1.3571 | 0.5943 |
